# Interactions between neutrophils and macrophages harboring gram-negative bacteria promote obesity-associated breast cancer

**DOI:** 10.1101/2024.08.08.607253

**Authors:** Sina T. Takle, Sturla Magnus Grøndal, Martin E. Lien, Priscilia Lianto, Wei Deng, Reidun Kristine Lillestøl, Per Lønning, James B. Lorens, Stian Knappskog, Nils Halberg

**Author notes:** Correspondence and requests for materials should be addressed to Nils Halberg.

## Abstract

Obesity promotes a more aggressive breast cancer phenotype. Through spatial and single-cell-based analysis of hormone receptor-negative breast cancers, we identify a subset of tumor-associated neutrophils (TANs) positive for granzyme B (GZMB) enriched in the tumor microenvironment of obese patients. In breast tumors evolved in obese environments, TANs are in proximity of M2 polarized macrophages containing lipopolysaccharides (LPS) from gram-negative bacteria. Pyroptosis of macrophages releases bacterial LPS, activating local GZMB^+^ TANs. This induces release of the S100 family member S100A8 that promotes tumor progression. In sum, we describe an obesity associated cellular network of cancer cells, neutrophils and M2 polarized macrophages that promotes tumor growth.

## INTRODUCTION

Obesity affects more than 2.5 billion adults globally (Collaborators, 2020) and is associated with a complex combination of comorbidities including insulin resistance (Unger and Scherer, 2010), leaky gut (Cani et al., 2007; Mishra et al., 2023), immunological dysregulation (Gregor and Hotamisligil, 2011), alteration of the commensal microbiome (Turnbaugh et al., 2006), as well as tumor risk and mortality (Bhaskaran et al., 2014; Calle et al., 2003). The cause of these comorbidities and especially the interactions between them are currently unknown.

Previous epidemiological analyses provide convincing evidence that obesity is associated with an increased risk of 13 cancers including postmenopausal (PM) breast cancer (Bhaskaran et al., 2014; Calle et al., 2003; Lohmann et al., 2021). The mechanisms of obesity-induced tumor initiation include cancer cell-autonomous epigenetic remodeling (Liu et al., 2022), lipid dependent signaling (Beyaz et al., 2016) and cancer cell-non-cell autonomous metabolic regulation of the tumor-immune interface (Tiwari et al., 2019). In addition to enhancing tumor initiation, the obese state is also linked to a more aggressive breast cancer (Chung et al., 2020; Incio et al., 2016a; Incio et al., 2016b; Petrelli et al., 2021), in part by negatively affecting the anti-tumor CD8 T-cell response (Ringel et al., 2020; Wang et al., 2019; Wogsland et al., 2021). At the metastatic site, obesity has been demonstrated to create more favorable conditions for cancer cell extravasation by inducing endothelial cell permeability governed by increased neutrophil NETosis (McDowell et al., 2021).

Bacteria are an integral part of mammalian physiology in health and disease and the microbiome is an enabling characteristic of cancer (Hanahan, 2022; Park et al., 2022). In recent years, emerging evidence have suggested that intracellular bacteria are present in multiple tumor types including colorectal, breast and lung cancers. Clinically, the abundance of such intracellular microbiomes is correlated with cancer subtypes (Nejman et al., 2020), cancer progression (Fu et al., 2022; Parhi et al., 2020), and therapy response (Bullman et al., 2017; Colbert et al., 2023; Geller et al., 2017). Despite the increased appreciation of obesity being a significant driver of multiple aggressive tumor phenotypes and associated with dysregulation of the microbiome, we still know very little of how the altered metabolic state in obese individuals affect the interactions between malignant, non-malignant cells, and their associated tumor microbiome. To fill this gap, we applied iterative imaging mass cytometry (IMC) analysis of treatment-naïve primary breast cancer tissue from PM obese and non-obese women to systemically define the consequence of obesity on the tumor microenvironment. Our findings demonstrate how obesity, through differential spatial interactions between the tumor microbiome and TANs, leads to enhanced primary tumor growth in PM hormone receptor negative (PM/ER^−^/PR^−^) breast cancer.

## RESULTS

### Systemic characterization of the tumor microenvironment in post-menopausal and hormone receptor negative breast cancer

To systemically compare the cellular composition and spatial interactions of the tumor ecosystem in PM/ER^−^/PR^−^ breast cancers from obese and non-obese patients, we collected paraffin embedded tissue sections from 12 patients with a body mass index (BMI) below or equal to 25 kg/m^2^ (termed non-obese) and from 22 patients with a BMI above 25 (termed obese; **Figure 1A**). We have previously shown that obese women in this cohort have shorter survival times than the non-obese group, indicative of a more aggressive cancer phenotype (Liu et al., 2022). All patients, independent of group, had similar age, hormone receptor status, tumor size, tumor stage (all stage III) and were collected pretreatment. We optimized a 36-marker antibody panel for IMC designed to broadly characterize the tumor microenvironment (**Figure 1B, Figure S1A**). Following data acquisition, single cells were segmented and signal spillover between neighboring cells was reduced by REDSEA (Bai et al., 2021) leading to the identification of a total of 197 747 cells. Using unsupervised (Levine et al., 2015) and supervised cell assignments, the cells were separated into 29 clusters including 12 cancer clusters and 17 non-cancer clusters (**Figure 1C**). Dimensionality reduction by Uniform Manifold Approximation and Projection (UMAP) yielded two main populations; one containing stromal- and one containing cancer cells (**Figure 1D**). Whereas the stromal cells overlapped between patients, the cancer cell clusters were highly patient specific reflecting expected between-patient tumor heterogeneity (**Figure S1B and C**).

**Figure 1.**
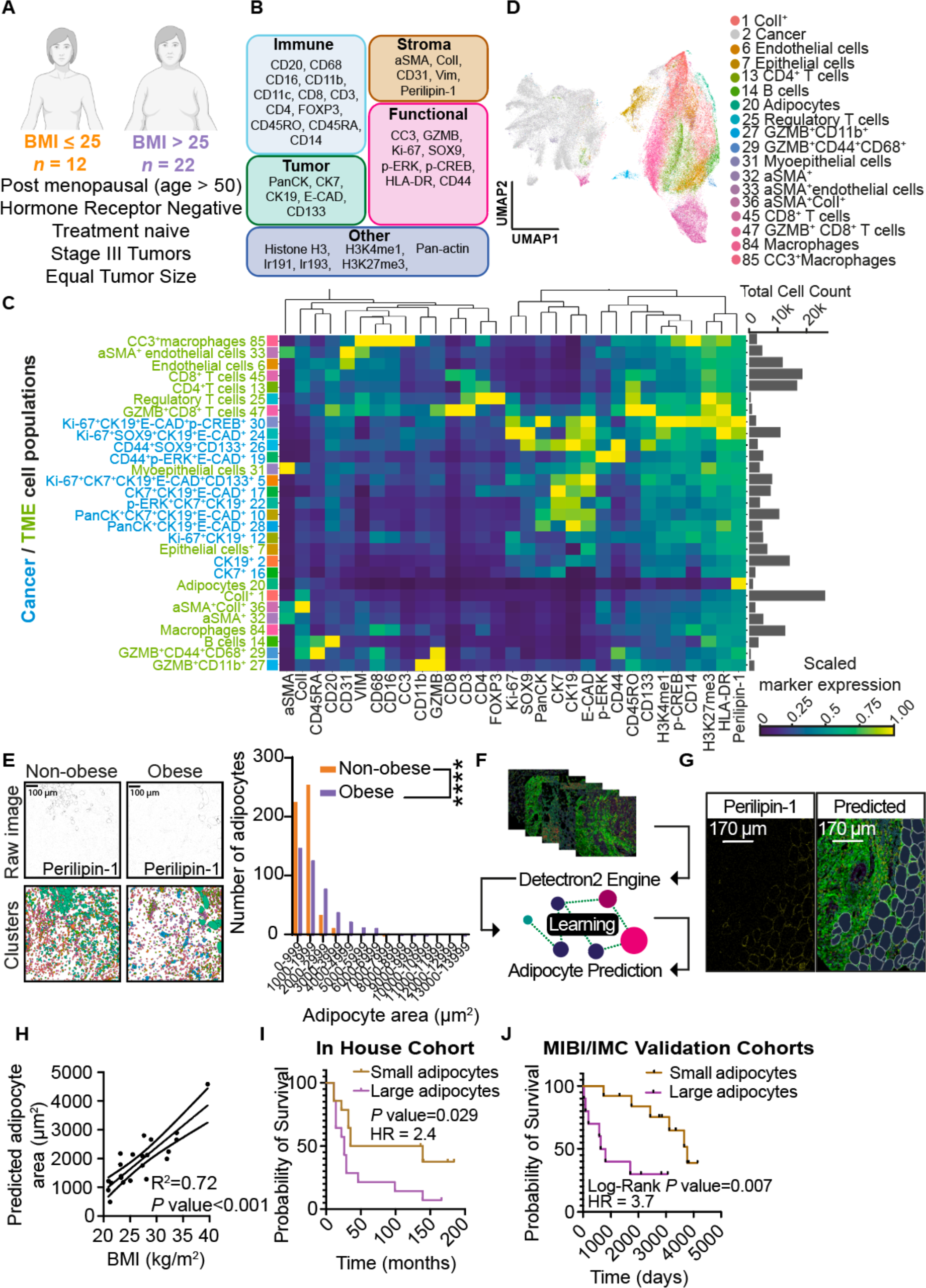
Systemic characterization of the tumor microenvironment in post-menopausal and hormone receptor negative breast cancer. (**A**) Overview of the patient cohort used for IMC analysis. Breast cancer tissue biopsies from non-obese (*n* = 12, BMI ≤ 25) and obese patients (*n* = 22, BMI > 25) were included. All patients were post-menopausal (patient age > 50 years), hormone receptor negative, had the same tumor size, stage III tumors and the biopsies were collected prior to treatment. (**B**) Markers included in the first IMC panel. (**C**) Heatmap of the arcsinh transformed (cofactor = 1 and addition of random noise (0-0.1)) mean marker expression of cell clusters generated by PhenoGraph (k=30) and manual curation for IMC panel 1. Each marker is range scaled to have a maximum value of 1. Cancer cell clusters are colored blue, while stromal cell clusters are colored green. Each cell cluster is assigned a color. Total cell count for each cell cluster is displayed. (**D**) UMAP plot displaying the identified cell clusters (in total 18 cell clusters) within the breast tumor microenvironment. (**E**) Representative images of adipocyte sizes within the breast cancer tissue of a non-obese (left) and an obese patient (right), displayed by Perilipin-1 signal (upper images) representing the adipocytes, and by adipocyte masks (lower images) colored according by cluster membership (green). Frequency distribution plot of the adipocyte area (µm^2^) of adipocytes (CL20) from obese (*n* = 442 cells, purple) and non-obese (*n* = 524 cells, orange) patients. **(F**) Illustration of a streamlined process for the prediction of adipocytes in IMC images utilizing the Detectron2 framework. (**G**) Side-by-side images show the results of Perilipin-1 staining in an IMC image, representing adipocyte borders versus predicted adipocyte borders generated using our computer-vision model. (**H)** Correlation between BMI and the predicted average area (µm^2^) of adipocytes per patient from the in-house dataset. (**I**) Kaplan-Meier plot comparing survival probability over time between patients from the in-house dataset categorized as either “Small adipocytes” (light brown, *n* = 14) or “Large adipocytes” (light purple, *n* = 14). Log-rank (Mantel-cox) *P* value and Hazard Ratio (HR) demonstrating the difference in survival between these two patient groups. (**J**) Kaplan-Meier plot for the validation cohorts from the IMC and MIBI datasets comparing the survival probability over time between patients categorized with “Small adipocytes” (light brown, *n* = 13) or “Large adipocytes’’ (light purple, *n* = 10). Shown is Log-rank (Mantel-cox) *P* value and Hazard Ratio (HR) displaying the difference in survival between these two patient groups. In (**E)** Mann-Whitney U test was performed to test for significant difference (\*\*\*\**P* value <0.0001).

BMI as a proxy for obesity and its related metabolic consequences is convenient but a suboptimal parameter that fails to incorporate factors such as fat-to-muscle mass ratios and adipose tissue distribution relevant to metabolic health. Conversely, the size of individual adipocytes has been reported to correlate to adiposity and inversely correlate to overall metabolic health (Laforest et al., 2017; Ryden et al., 2014). We therefore used the spatial IMC data to quantify the adipocyte (CL20) size distribution between the obese and non-obese patient groups. The adipocytes of the high BMI group were larger, consistent with obesity (**Figure 1E**). Next, we reasoned that the unique morphology of adipocytes including large cell size, elongated nucleus, and lack of stain due to removal of lipid droplet during tissue preparation could be used to train a pattern detection algorithm to identify adipocytes in a panel-agnostic fashion. As BMI, or other measures of adiposity, are rarely available in large, published datasets and consortia, adipocyte size measurements in publicly available tumor images could enable validation in independent datasets. We developed a robust computer vision model using the Detectron2 framework (Wu et al., 2019). The model was trained on annotated dataset containing known adipocytes (CL20) and subsequently validated on images that were not part of the training dataset (**Figure 1F**). With this setup the model achieved an F1 score of 82% and an average precision at IoU (0.50:0.95) of 82.2% ((Muller et al., 2022); **Figure S1D**). The predicted adipocyte size correlated with BMI (R^2^=0.72, *P* value <0.001; **Figure 1H**) and large adipocyte size were associated to shorter survival rates (**Figure 1I**). We then identified imaging datasets based on both MIBI (Keren et al., 2018) and IMC (Jackson et al., 2020) containing patient cohorts closely mimicking our original cohort parameters (PM/ER^−^/PR^−^ breast cancer patients) and applied our computer vision model. While the number of observations for each cohort was limited, combining the data, we found that individuals with large adipocytes had a significantly worse prognosis than those with small adipocytes (*P* value = 0.007; **Figure 1J**). Further, when applying a multivariate model including datasets as a covariate, this association remained significant (*P* value = 0.017) confirming that obesity and obesity-related unhealthy metabolic health is correlated to a more aggressive breast cancer phenotype across patient cohorts.

### The obese tumor microenvironment is characterized by an increased abundance of neutrophils and CD4^+^ T-cells and fewer endothelial cells

The tumor microenvironment is characterized by cellular neighborhoods comprising divergent cell types (Schurch et al., 2020). To determine if obesity affects this level of tumor architecture, we randomly divided tissue sections into 5 phenotypically similar areas based on their cell type composition (Stoltzfus et al., 2020). This resulted in 5 regions enriched in 1) immune cells; 2) extracellular matrix (ECM) associated cells; 3) cancer cells, 4) ECM, and 5) the tumor-immune cell boundary (**Figure 2A**, **Figure S2A and B**). At this level, the tumors evolved in obese and non-obese patients were similar (**Figure 2B**). We next asked if the proportion of each cell cluster within regions or entire ROI differed between tumors evolved in obese compared to non-obese patients. Given the patient tumor cell heterogeneity, we focused this analysis on non-cancer cell clusters. This revealed that CL13 (CD4^+^ T cells) and CL27 (GZMB^+^ CD11b^+^ cells) cells were enriched in the obese state in region 1 and 2, respectively. CL6 (CD31^+^ endothelial cells) were reduced specifically in the cancer region in the obese patients (**Figure 2C and D**). This unbiased approach firstly provides clinical relevance to previous murine findings that tumors grown in obese mice display reduced amount of blood vessels and increased abundance of CD4 memory T-cells (Incio et al., 2016a; Incio et al., 2016b; Wang et al., 2019). The cell population most affected by obesity was CL27, that were more abundant both in region 2 (3.80-fold; *P* value = 0.0197) as well as within all regions combined (2.55-fold; *P* value = 0.0381). Based on the marker expression of CL27 (**Figure 1C and 2E**) this cell cluster potentially contained several cell types, including macrophages, neutrophils, or NK cells. We therefore performed immunofluorescence of TMA serial sections. This analysis demonstrated that the GZMB^+^ CD11b^+^ cells also expressed myeloperoxidase (MPO) but not CD56 and CD68, suggesting that these cells are TANs (**Figure 2F, Figure S2C**).

**Figure 2.**
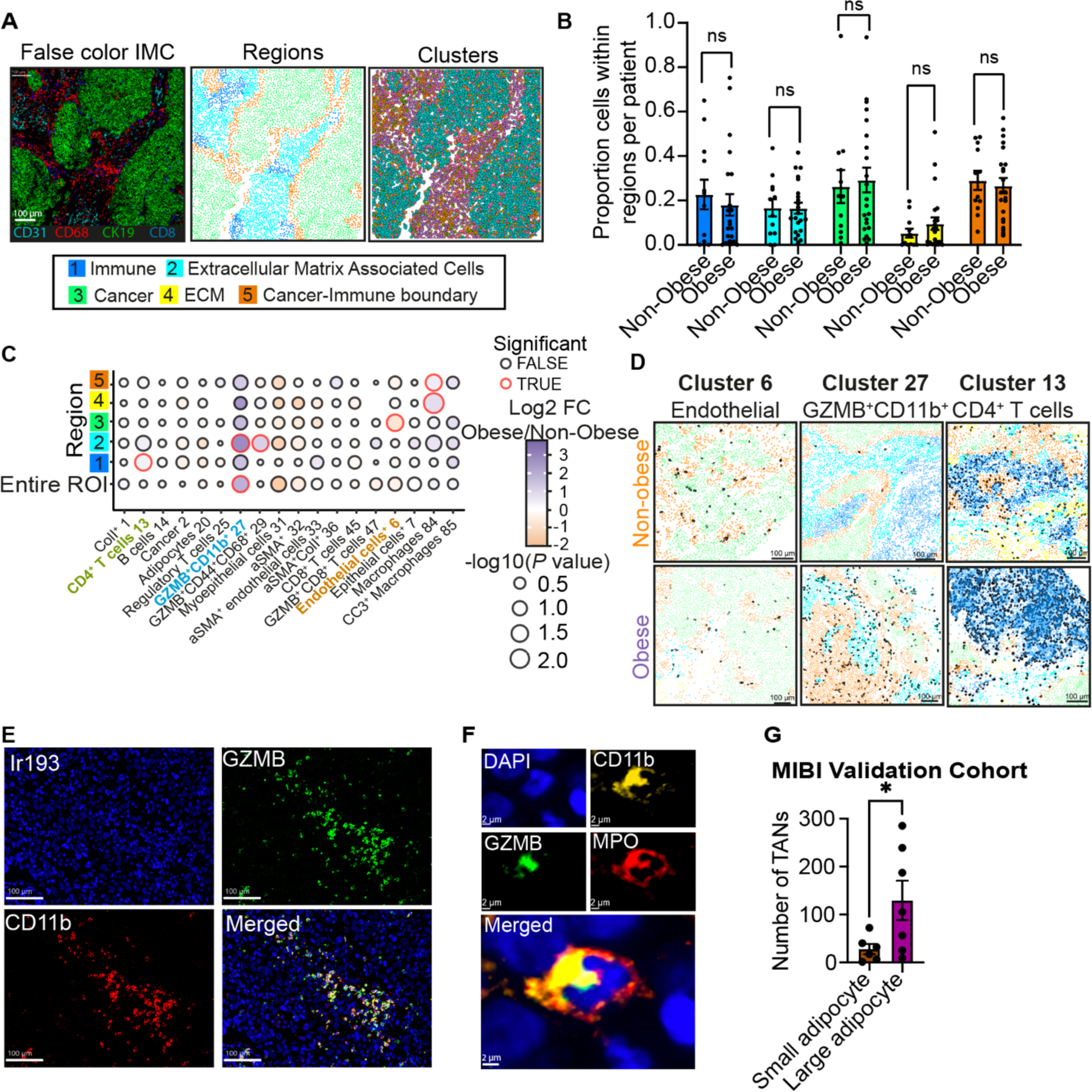
The Obese Tumor Microenvironment is Characterized by an Increased Abundance of Neutrophils and CD4^+^ T-cells and reduced Endothelial Cells. (**A**) Five regions within the breast TME were generated based on similar cell cluster composition. IMC image colored by markers representing cell types found in different regions CD31 (cyan), CD68 (red), CK19 (green), and CD8 (blue) (left image), region (middle image), and PhenoGraph clusters (right image). (**B**) Proportion of cells within each region (1-5) in obese (*n* = 22) and non-obese (*n* = 12) patients. Each region is labeled with an individual color, region 1 (blue), region 2 (cyan), region 3 (green), region 4 (yellow) and region 5 (orange). Mean with SEM displayed in the bar plot. (**C**) Difference in cell cluster proportion within each region between obese and non-obese patients. Bubble plot visualizes the difference in proportion of each cell cluster (x-axis) within each region (y-axis) between obese and non-obese patients. The bubble color represents the log2FC of obese versus non-obese patients of the cell cluster proportion within each region. Purple represents an increased cell cluster proportion in obese patients, while orange represents an increased cell cluster proportion in non-obese patients. The bubble size increased with lower *P* value in the form of -log10(*P* value). Significant difference (*P* value < 0.05) in cell cluster proportion is highlighted in red. Missing bubbles are due to too few observations for a specific cell cluster in one or both groups. (**D**) Representative images of a specific cell cluster’s spatial distribution within regions. The cells of the specific cell cluster of interest (cluster 6, cluster 27 and cluster 13) are visualized in black, while the regions are colored as in (A). There is one representative image for each condition. (**E**) Representative IMC images of cells expressing GZMB (green) and CD11b within an ROI. Iridium 193 (Ir193) display nuclei in blue. (**F**) Representative image of a GZMB^+^ neutrophil. Immunofluorescence (IF)-staining of a GZMB^+^ neutrophil from a breast cancer patient TMA core, demonstrating that GZMB^+^ (green) neutrophils also express CD11b (yellow) and MPO (red). (**G**) Quantification of tumor infiltrating neutrophils (TANs) in patients with large adipocytes compared to non-obese from a MIBI validation cohort. In (**B**) Mann-Whitney U test was performed for statistical testing (* *P* value < 0.05). In (**C**) to test for statistical significance in the proportion of cells in a specific region between non-obese and obese patients, Welch’s two-sample t test was utilized. To test for statistical differences in cell cluster proportion from the entire core, Kruskal-Wallis rank sum test was performed. In (**G**) statistical testing was done using two-tailed Students t-test.

To independently validate that breast tumors grown in metabolically unhealthy patients have increased abundance of TANs, we stratified patients based on adipocyte size (**Figure 1I and J**). We used the MIBI primary breast cancer dataset derived by Keren et al., (Keren et al., 2018) which included the neutrophil marker MPO. Consistent with our data, PM/ER^−^/PR^−^ patients with enlarged mammary adipocytes displayed increased number of TANs (**Figure 2G**). These findings confirm that PM/ER^−^/PR^−^ breast cancers from obese patients are characterized by an increased abundance of TANs and CD4^+^ T-cells and reduced angiogenesis.

### Obesity promotes the interaction between GZMB^+^ tumor-associated neutrophils, gram-negative bacteria, and M2-polarized macrophages

Neutrophils normally lack granzyme expression, but GZMB^+^ neutrophils have been reported to interact with lipopolysaccharides (LPS) positive gram-negative bacteria (Martin et al., 2018) and gram-negative *Mycobacterium tuberculosis* (Mattila et al., 2015). To test if LPS abundance is associated with GZMB^+^ TANs in CL27, we evaluated tissue sections from PM/ER^−^/PR^−^ breast cancer patients with antibodies targeting LPS (**Figure 3A**). Patient tumor LPS levels correlated with the proportion of TANs (CL27; R^2^ = 0.72, *P* value = 7.01 × 10^−5^) and CL85 (R^2^ = 0.64; *P* value = 7.06 × 10^−4^; **Figure 3B**). CL85 cells were defined by positivity for CD68, CD11b, CD14, CD16, and cleaved caspase 3 (CC3) suggesting these are macrophages undergoing CC3-dependent cell death (**Figure 1C**). CC3 is involved in multiple forms of cell death including apoptosis and pyroptosis (Jiang et al., 2020). Immunostaining of 100 CC3^+^ macrophages, showed consistent co-staining of gasdermin E, indicating that these cells are undergoing pyroptosis, a lytic form of cell death that exposes intracellular bacteria (**Figure S3A**).

**Figure 3.**
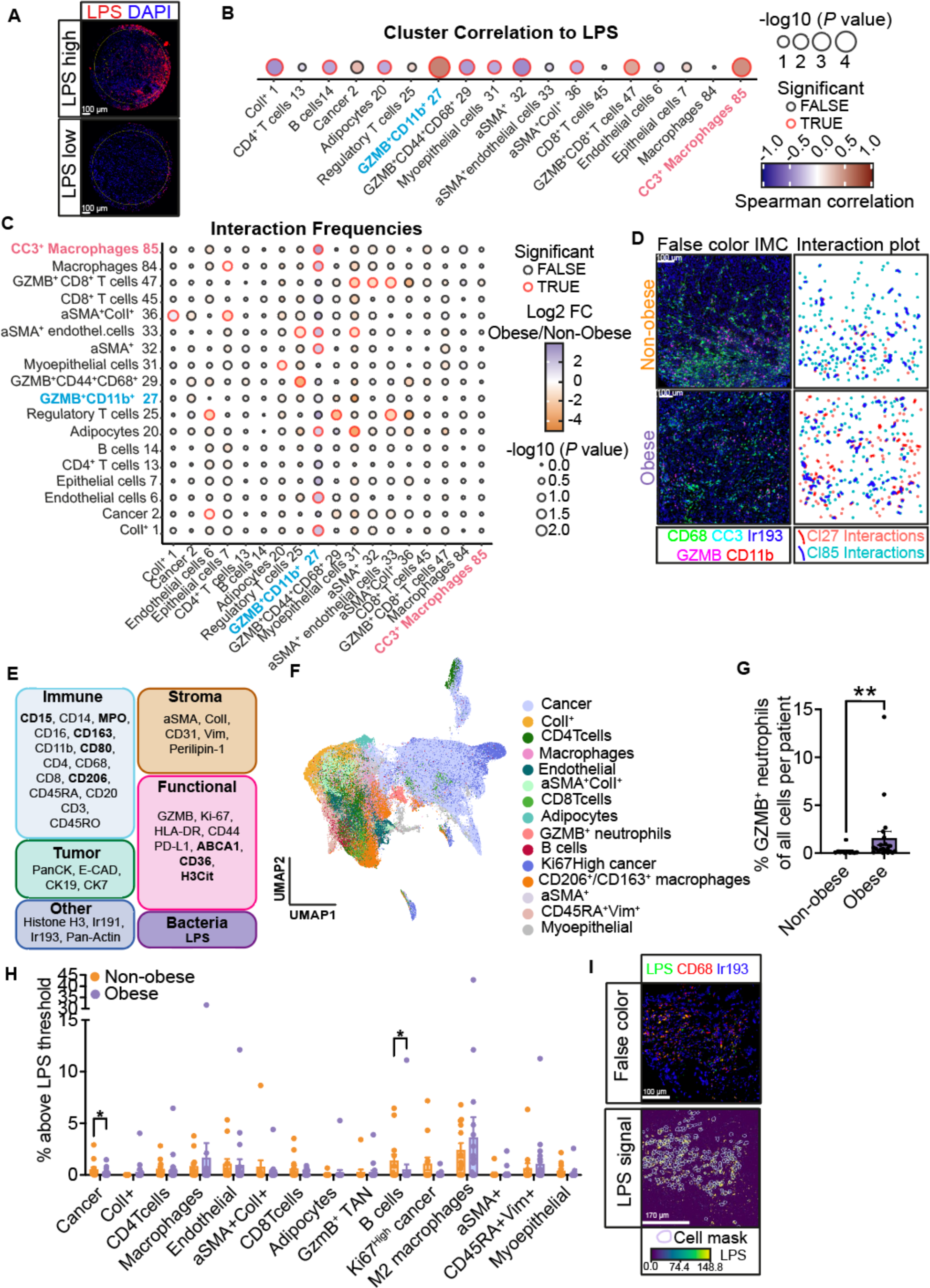
The Obese Environment Governs the Interaction Between GZMB^+^ Tumor Associated Neutrophils, Gram-negative Bacteria, and M2-polarized macrophages. (**A**) Representative IF images of human breast cancer TMA core with high LPS signal (upper image), and low LPS signal (lower image). LPS and DAPI are visualized in red and blue, respectively. The annotated area (yellow circle) was used for LPS signal quantification. (**B**) Correlation of IMC panel 1 cell cluster abundance to mean IF LPS signal intensity for each patient (*n* = 25). The correlation is displayed in a bubble plot, where the color of the bubbles represents the Spearman correlation (−1 to 1, blue and red, respectively) for the abundance of each cell cluster to mean LPS signal intensity. Bubble size increases with significance (-log10 of *P* value). Significant correlations are highlighted in red. (**C**) Interaction frequency plot of cell cluster interactions displaying whether a cell cluster (row) has more neighbors of another cell cluster (column) in obese than non-obese. The bubbles are colored by the log2FC between mean interaction counts in obese versus non-obese to visualize an increase in obese (purple) or non-obese (orange). Bubble size represents a significant difference in mean interactions between obese and non-obese patients (-log10(*P* value)). The rows display which cell cluster that are surrounding the cell type of interest. The columns demonstrate which cell cluster the cell type of interest is surrounding. (**D**) Spatial interaction between cells from cluster 27 and cluster 85. False color IMC images (right images) visualizing the CD68 (green) and CC3 (cyan) expression within an entire representative non-obese (orange) and obese (purple) patient core, representing CC3^+^ macrophages (cluster 85) and the GZMB (magenta) and CD11b (red) expression within a representative non-obese and obese patient core, representing GZMB^+^CD11b^+^ cells (Cluster 27). The spatial interaction (left images) between Cluster 27 (red) and Cluster 85 (blue) were visualized for the same patient cores. Interactions were visualized by edges, where red represents Cluster 27 interactions while blue displays Cluster 85 cell interaction. (**E**) Overview of the markers included in the second IMC panel. New markers added in the IMC panel 2, primarily targeting neutrophils, macrophages, and bacteria, compared to the first IMC panel in bold (**F**) UMAP plot displaying the identified cell clusters (total 15 cell clusters) utilizing the markers in the second IMC panel. Each cell is colored by cell cluster membership. (**G**) Bar plot of the percentage of GZMB^+^ neutrophils in non-obese (*n* = 11, orange) and obese (*n* = 21, purple) patients (*P* value = 0.0048) displaying mean with SEM. (**H**) Bar plot displaying the percentage of LPS^+^ cells for each cell cluster, found with IMC panel 2. An LPS^+^ cell was characterized as having an LPS signal above the LPS threshold which was set to the 99^th^ percentile of LPS signal for all cells. There was significantly a higher percentage of LPS^+^ Cancer cells (*P* value = 0.0135) in the non-obese compared to the obese patients. The bar plots are displayed with mean with SEM. (**I**) Spatial visualization of CD206^+^/ CD163^+^ macrophages colocalization with LPS signal displayed with false color image (upper image) of LPS (green), CD68 (red) and Ir193 (blue), and LPS signal (lower image) overlaid with cell masks (light purple) from the CD206^+^/CD163^+^ macrophage cluster. For (**C**) two-tailed Mann-Whitney U test was used to test for significant difference between the mean interaction count between obese and non-obese patients for each cell cluster. For (**G**) two-tailed Mann-Whitney U test was used to test for statistical significance. For (**H**) statistical testing comparing the percentage of cells above the LPS threshold was performed for all cell clusters using two-tailed Mann-Whitney U test.

Given that both CL27 and CL85 were associated with LPS abundance, we next sought to define their spatial relationship. An unbiased neighborhood analysis demonstrated that CL27 cells interact with CL85 cells at significantly higher frequency in the obese compared to non-obese patient samples (mean interaction count for obese patients = 0.174, mean interaction counts for non-obese patients = 0.012, log2FC = 3.820, *P* value = 0.0398); **Figure 3C and 3D**).

To better understand the nature of the interplay between the obese environment, TAŃs, dying macrophages, and the tumor microbiome, we generated a second IMC antibody panel including markers for macrophage differentiation, neutrophil function and LPS (**Figure 3E**). Following cell segmentation and cluster annotation, this analysis identified 207 956 individual cells divided over 15 clusters (**Figure 3F, Figure S3B,C**). Consistent with the first IMC analysis, the obese environment was characterized by increased proportion of GZMB^+^ MPO^+^ TANs (**Figure 3G**). Neutrophils can undergo NETosis, a form of cell death that releases chromatin “nets” containing bactericidal proteins. These structures have been independently linked to the obese phenotype and to primary tumor growth (Adrover et al., 2023; Shaul and Fridlender, 2019). However, when quantified as signal overlap between citrullinated histone H3 and MPO we found only sparse NET formation in both obese (6 out of 22 patients) and non-obese (1 in 13 patients) patients, suggesting that NETosis is not associated with obesity-induced primary tumor progression (*P* value = 0.22; **Figure S3D**).

We next surveyed all cell clusters for the presence of gram-negative bacteria through LPS positive signal. Overall, we found no significant difference in the frequency of LPS^+^ cells between obese and non-obese patients (0.651% in obese vs 1.038 % in non-obese). A cluster specific analysis demonstrated that the presence of LPS were most abundant in M2-polarized macrophages (3.4% of M2-polarized macrophages were LPS^+^; **Figure 3H and 3I**). While LPS positivity in M2-polarized macrophages was similar between obese and non-obese patients, it was more frequent in the non-obese setting for CK19^+^ cancer cells and B cells (**Figure 3H**).

Thus, iterative IMC and immunofluorescence analysis of the tumor microenvironment reveals that the obese state leads to increased cellular interactions between GZMB^+^ TANs and dying macrophages and the presence of both cell types positively correlates with the LPS content in the tumor. Importantly, LPS levels are similar in magnitude and cell-specificity between obese and non-obese breast cancer patients, suggesting that the differential tumor properties are not caused by the presence of LPS itself, but rather the immune response to the LPS^+^ cells governed by GZMB^+^ neutrophils.

### The tumor microbiome in breast tumors is not altered in obese environments

To unbiasedly extend our analysis of the obese tumor environment to all bacteria, we next performed 16S sequencing of frozen tumors from the same sample set of PM/ER^−^/PR^−^ breast cancer patients. While we cannot control potential bacteria contamination from tissue removal in this retrospective cohort, we were able to robustly detect 39 bacterial genera (**Figure 4A**). Across all patients, we demonstrate that a higher bacterial diversity defined by the Shannon Index is associated with better relapse free survival outcomes (**Figure 4B**). When stratifying patients according to BMI, we find that higher diversity is trending towards improved survival in both the obese group and non-obese groups (**Figure 4C** and **Figure S4A)**. Overall, we do not find any overall differences in the tumor microbiota composition or diversity between obese and non-obese PM ER-/PR-breast cancer patients. This suggests that the obesity-induced tumor growth promotes a unique TAN response to the presence of LPS, rather than the presence/absence of specific bacteria.

**Figure 4.**
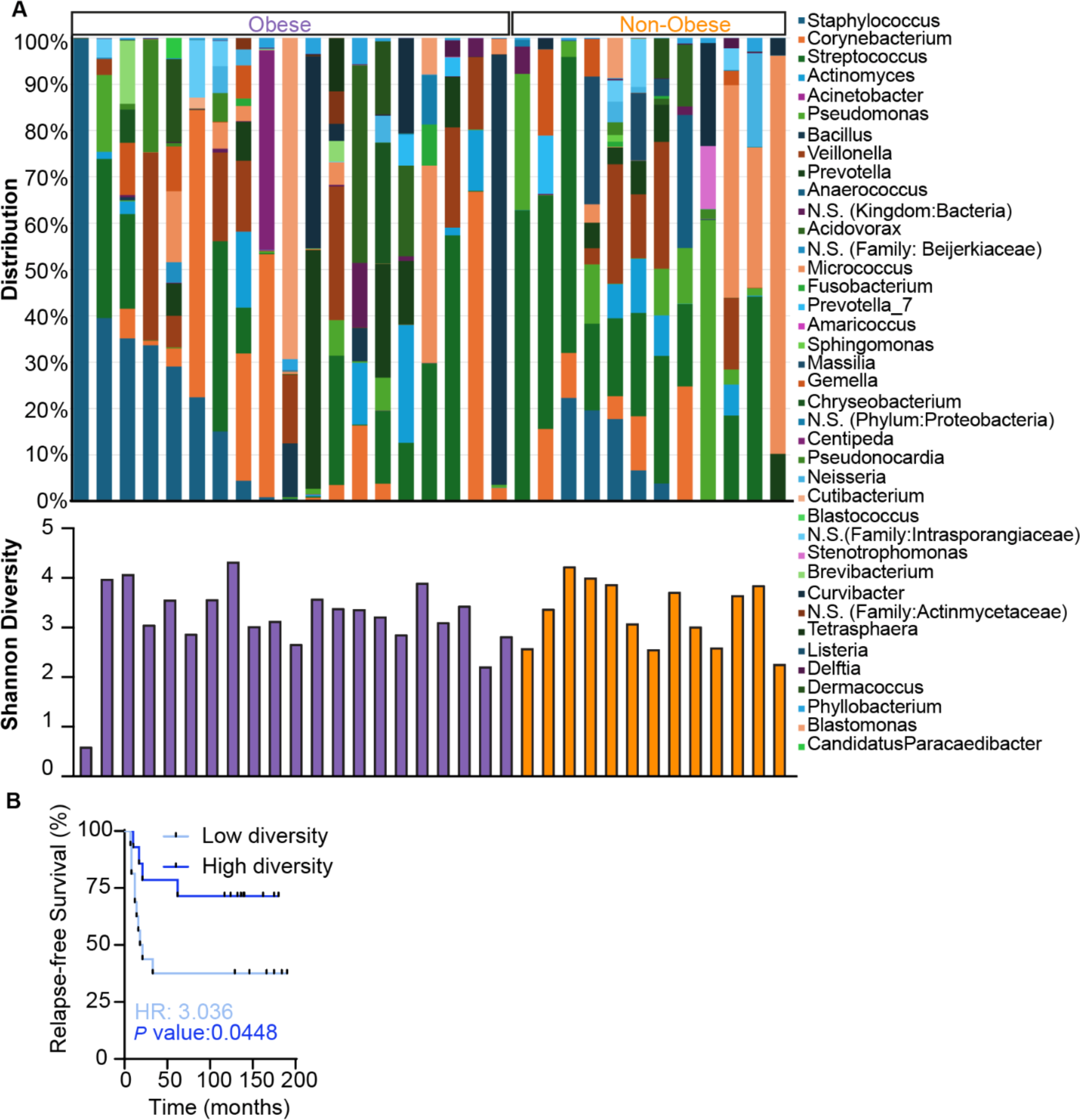
The tumor microbiome in breast cancer is not affected by obese environments. (**A**) *Upper panel*, frequency distribution of the 39 identified bacteria (genus) identified by 16S sequencing across all included patients (N.S.; not specified on the genus level). *Lower panel*, the Shannon diversity score for each patient. (**B**) Kaplan-Meier plot comparing relapse free survival between patients from the in-house dataset categorized as either high or low tumor microbiota diversity based on a median split by the Shannon diversity score including all patients. Log-rank (Mantel-cox) *P* value and Hazard Ratio (HR) were used to determine the difference in survival between these two patient groups.

### Tumor-associated neutrophils of the obese tumor microenvironment are immature and respond differentially to LPS than in non-obese states

We next sought to better understand the mechanistic basis of how the obese environment affects the TAN abundance and their response to LPS stimulation. Mice were fed a high fat or regular diet and syngeneic PyMT mammary tumor cells were implanted. As expected, the high fat diet fed mice were obese as determined by larger fat pads, body weight, liver steatosis and tumors grew larger (**Figure 5A-D**). TANs were then isolated from the primary tumors and subjected to RNASeq analysis (**Figure 5E**). Principal component analysis demonstrated that TANs isolated from tumors grown in obese mice were phenotypically distinct from those in non-obese mice (**Figure 5F**). GSEA analysis revealed an enrichment in oxidative phosphorylation in obese mice (NES = 1.69, FDR q-val = 0.002) and EMT (NES =-1.94, FDR *q* value = 0.000) enriched in the lean mice (**Figure 5G**). To phenotypically confirm the transcriptional data, we isolated TANs from PyMT tumors grown in obese and non-obese mice and conducted metabolic profiling and tracing of radio-labelled glucose and palmitate into CO_2_. Both assays demonstrated that TANs isolated from an obese environment rely more on oxidative phosphorylation irrespective of substrate use (**Figure 5H,I,J**).

**Figure 5.**
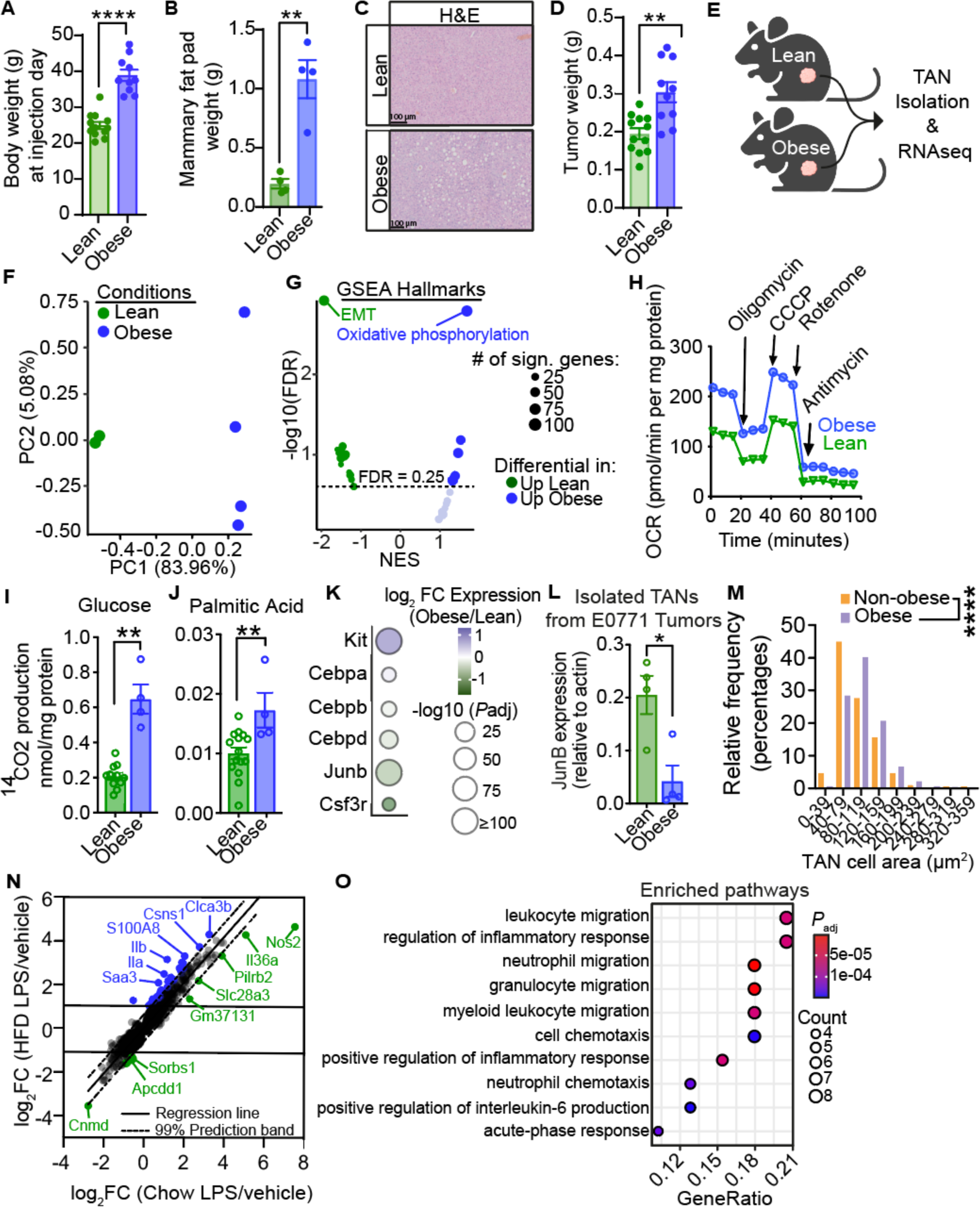
Tumor-associated neutrophils of the obese tumor microenvironment are immature and respond differentially to LPS than in non-obese states. (**A**) Body weight (g) of obese (*n* = 10, blue) and lean (*n* = 12, green) female C57BL/6 mice at the day of injection of 1000 PyMT cells orthotopically into the 4^th^ inguinal mammary fat pad. (**B, C**) The weight of the 4^th^ inguinal mammary fat pad (B) and liver H&E histology (C) from obese (*n* = 4, blue) (fed HFD for 12 weeks before implantation) and lean (*n* = 4, green) (fed Chow for 12 weeks before implantation) female C57BL/6 mice 29 days after injection (*P* value = 0.0095). (**D**) Weight of tumors derived from PyMT cells developed in the 4^th^ inguinal mammary fat pad in obese (*n* = 10, blue) and lean (*n* = 12) C57BL/6 mice for 28 and 30 days (*P* value = 0.0011). (**E**) Female C57BL/6 mice fed HFD (obese), or Chow (lean) for 18 weeks were orthotopically injected with PyMT cells into the mammary fat pad. Neutrophils were isolated from dissociated tumors. Neutrophils used for RNA-seq were cultured with or without LPS before RNA-extraction and RNA-seq. (**F**) PCA plot of RNA-seq counts from PyMT tumor derived neutrophils from obese (blue) and lean (green) female C57BL/6 mice. (**G**) GSEA hallmark analysis. Volcano plot displaying the hallmark gene sets that are differentially expressed in tumor derived neutrophils from obese (blue) and lean (green) mice. A significant difference was set to a false discovery rate (FDR) equal to 0.25. NES = normalized enrichment score. (**H**) Oxygen consumption rate (OCR) measured for PyMT tumor derived neutrophils from obese (blue) and lean (green) mice fed HFD or Chow for 11 weeks before implantation. (**I**) Glucose and (**J**) palmitic acid oxidation of neutrophils derived from PyMT tumors from obese (blue) and lean (green) mice fed HFD or Chow for 10 weeks. (**K**) Bubble plot displaying the difference in gene expression of genes associated with neutrophil maturity, either immature (Kit, Cebpa) or mature (Cebpb, Cebpd, Junb and Csf3r) from PyMT tumor derived neutrophils from obese (blue) and lean (green) mice. Bubble size represents the significant difference between obese and lean mice. (**L**) Junb expression relative to actin in EO771 tumor derived neutrophils grown in lean (*n* = 4, green) and obese (*n* = 4, blue) mice (*P* value = 0.0133) fed HFD or Chow for 12 weeks before implantation. Mean with SEM is denoted in the bar plot. (**M**) Frequency distribution plot of TAN cell area (µm^2^) from obese (*n* = 1669 cells, purple) and non-obese (*n* = 191 cells, orange) patients (*P* value < 0.0001). The TAN cell area was found utilizing the cell area of CL27 in IMC panel 1. (**N**) Comparison of LPS-stimulated genes in TANs isolated from obese and non-obese mice. Highlighted in blue and green are genes that falls outside the 99% prediction bands. (**O**) GO: BP enrichment analysis of the genes that were specifically induced by LPS in the neutrophils isolated from obese tumors. For (**A**), (**D**) and (**L**), significant differences were tested using t-test (\*\*\*\**P* value <0.0001). For (**B**) statistical testing was performed by Welch’s t-test. For (**I**) and (**J**) statistical testing was done by students t-test. For (**M**), statistical testing was conducted using two-tailed Mann-Whitney U test (\*\*\*\**P* value < 0.0001). For the bar plot of (**A, B, D, I, J** and **L**) mean with SEM is displayed.

TANs are short-lived and metabolically active cells and increased reliance on oxidative phosphorylation is characteristic of immature neutrophils (iTANs; (Jeon et al., 2020)) that are generally associated with a pro-tumor phenotype (i.e., decreased phagocytosis and oxidative bursts; (Sagiv et al., 2015)). In addition to these functional assays, we determined the expression of differentiation genes (including c-Kit and CEBPBA) for immature neutrophils were increased while markers of mature neutrophils (CEBPB/D, JunB and Csf3r) were all decreased (**Figure 5K**). To exclude the possibility that this neutrophil phenotype being PyMT tumor specific, we similarly evaluated the syngeneic E0771 mammary carcinoma model (**Figure S5A,B**). Quantitative PCR of isolated TANs from E0771 tumors grown in obese mice, confirmed loss of mature neutrophil marker JunB (**Figure 5L**). Collectively, this transcriptional data suggests that TANs from tumors formed in obese mice remain immature. Morphologically, immature neutrophils have been suggested to be larger than fully matured neutrophils (Sagiv et al., 2015). Consistently, TANs from the obese patient were larger than in non-obese (**Figure 5M**). Given the spatial relationship between LPS^+^ dying macrophages and GZMB^+^ neutrophils in the tumor microenvironment, we next questioned if the obese TAN phenotype is associated with a differential response to LPS. We thus treated isolated TANs *in vitro* with LPS and repeated the RNAseq. To extract genes that were differentially affected by LPS in TANs isolated from obese and lean mice, we correlated the log2 fold change between control and LPS stimulated cells in both conditions and extracted genes who were differentially affect by LPS between obese and lean mice (**Figure 5N**). This identified 43 genes that were more stimulated (highlighted in blue) and 16 genes that were more repressed (highlighted in green) upon LPS stimulation in the TANs isolated from obese tumor microenvironments. This included genes enriched in pathways related to neutrophil migration and cell chemotaxis (**Figure 5O**) suggesting a link to the increased number of neutrophils in the tumors from obese breast cancer patients.

In sum, neutrophils from tumors evolved in the obese setting are immature and react differentially to LPS stimulation to activate cellular migration pathways.

### LPS-dependent iTAN secretion of S100A8 leads to enhanced tumor aggressiveness

The shorter survival of obese PM breast cancer patients reflects a more aggressive cancer phenotype. We hypothesized that an altered neutrophil response to LPS could drive this through paracrine signaling. To identify potential drivers of such signaling events, we reexamined the TAN transcriptional data and searched for genes coding for secreted proteins who was relatively unaffected by LPS stimulation in the lean setting while being induced in the obese. This analysis highlighted S100 calcium-binding protein A8 (S100A8) and Casein Alpha S1 (Csn1s1) as being exclusively expressed by LPS in TANs from obese environments (**Figure 6A**). Importantly, this obesity-dependent LPS stimulation was consistent in neutrophils isolated from E0771 tumors (**Figure 6B**) To support this mouse-based analysis and determine its relevance for human disease, we analyzed survival rates in obese and non-obese patients stratified by their tumor S100A8 and Csn1s1 expression levels in our in-house patient cohort. Whereas no differences were observed when stratifying the patients based on Csn1s1expression, we found that high expression of S100A8 was associated with shorter survival exclusively in the obese state (**Figure 6C, Figure S6A-C**). To further validate these findings, we extracted known BMI from the Metabric dataset (Nguyen et al., 2023) and correlated patient survival rates to expression of S100A8 in PM/ER^−^/PR^−^negative patients. Consistent with our in-house dataset, high expression of S100A8 was significantly associated with shorter survival rates in obese breast cancer patients (**Figure 6D**). Further, patients with high expression of S100A8 also had larger tumors at diagnosis (**Figure 6E**) consistent with a more aggressive cancer phenotype. Finally, given that S100A8 is highly expressed by neutrophils (Ryckman et al., 2003), we extracted genes whose expression positively correlated to that of S100A8 and identified associated expression programs reflecting the biological function of S100A8. Interestingly, this unbiased analysis of the Metabric dataset, highlight multiple pathways involved in response to bacteria, again linking the obesity-dependent iTAN phenotype to tumor bacteria (**Figure 6F**). Combined, this data demonstrates that tumors growing in obese environments are associated with iTANs that, upon LPS stimulation, leads to increased production of the cytokine S100A8. And further, suggests that such iTAN-derived S100A8 is a mediator of a more aggressive tumor type specifically in obese breast cancer patients. We next sought to determine the mechanism by which S100A8 could be involved in tumor progression. S100A8 is an ill-defined pro-inflammatory cytokine that can act as a homodimer or in heterodimer together with S100A9 (also called calprotectin; (Gebhardt et al., 2006)). Two independent breast cancer cell lines treated with recombinant S100A8 homodimer or S100A8/S100A9 heterodimer displayed enhanced proliferation and tumorsphere formation (**Figure 6G-J**). This supports that S100A8 released by the iTANs could be a driver of obesity-induced tumor growth.

**Figure 6.**
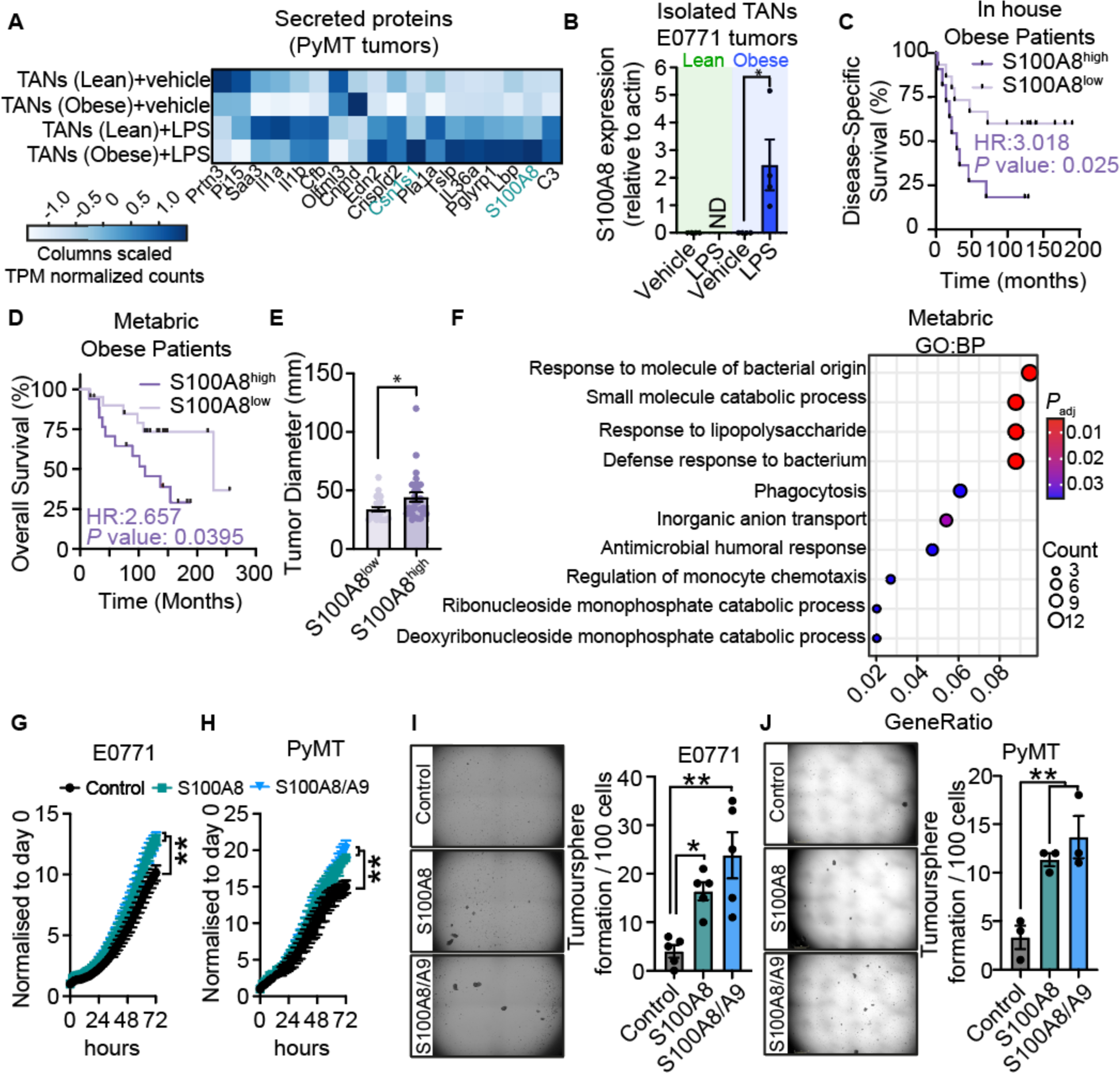
LPS-dependent iTAN secretion of S100A8 leads to enhanced tumor aggressiveness. (A) Genes identified in Fig 5N as being exclusively induced by LPS in TAN from obese mice containing a signal peptide sequence were extracted. Expression of these genes are displayed in a heatmap with mean TPM normalized count values scaled for each gene across each sample group. (B) S100A8 expression relative to actin of EO771 tumor derived neutrophils from obese (blue) or lean (green) mice fed HFD or Chow for 12 weeks before implantation treated with LPS or vehicle (*P* value = 0.037). ND = not detected. (**C**) Kaplan-Meier plot displaying Disease-Specific Survival of obese (BMI > 25 kg/m^2^) ER^−^PR^−^ postmenopausal (50 > years) breast cancer patients from the in-house data set. The patients were separated based on S100A8 expression of both obese and lean (BMI ≤ 25 kg/m^2^) patients (median = 17.572), where the obese patient separation was set as S100A8^high^ > median (dark purple, *n* = 11) and S100A8^low^ ≤ median (light purple, *n* = 15). Log-rank (Mantel-cox) test *P* value and Hazard ratio (HR) (logrank) are displayed. (**D**) Kaplan-Meier plot displaying Overall Survival of obese (BMI> 25) ER-PR-post-menopausal (50> years) breast cancer patients from the METABRIC cohort. The patients were separated based on normalized S100A8 expression of obese and lean patients (median = 0.916), where the obese patient separation was set as S100A8^high^ (dark purple, *n* = 17), and S100A8^low^ (light purple, *n* = 22). Log-rank (Mantel-cox) test *P* value and Hazard ratio (HR) are displayed. (**E**) Tumor diameter (mm) of obese (BMI > 25) ER-PR-postmenopausal (50 > years) breast cancer patients from the METABRIC cohort. The patients were separated based on normalized S100A8 expression of obese and lean patients (median = 0.916), where the obese patient separation was set as S100A8^high^ (dark purple, *n* = 28), and S100A8^low^ (light purple, *n* = 24). (**F**) GO:BP of genes correlating (R^2^>0.4) with S100A8 specifically in obese PM/ER^−^/PR^−^ patients in the Metabric dataset. Proliferation assay of EO771 (**G**) and PyMT (**H**) cells cultured with recombinant S100A8 (5 µg/ml, green) or S100A8/S100A9 (5 µg/ml, blue). (**I**) Tumorsphere assay of EO771 and (**J**) PyMT cells treated with recombinant protein of S100A8 (5 µg/ml, green) or S100A8/A9 (5 µg/ml, blue). For (**B,E,I,J**) a Student’s t-test was used to test for statistical difference. For (**G**) and (**H**) significant differences were tested using Ordinary one-way ANOVA with Holm-Šídák’s multiple comparisons test. All bar graphs display the mean with SEM.

## DISCUSSION

Our findings establish a cellular interaction network between bacteria-containing M2-polarized macrophages, immature neutrophils and tumor cells that drives obesity-induced post-menopausal breast cancer. We demonstrate that the obese environment uniquely supports immature TANs that upon stimulation with LPS, releases the pro-tumorigenic cytokine S100A8, driving acquisition of aggressive cancer cell phenotypes (**Figure 7**).

**Figure 7.**
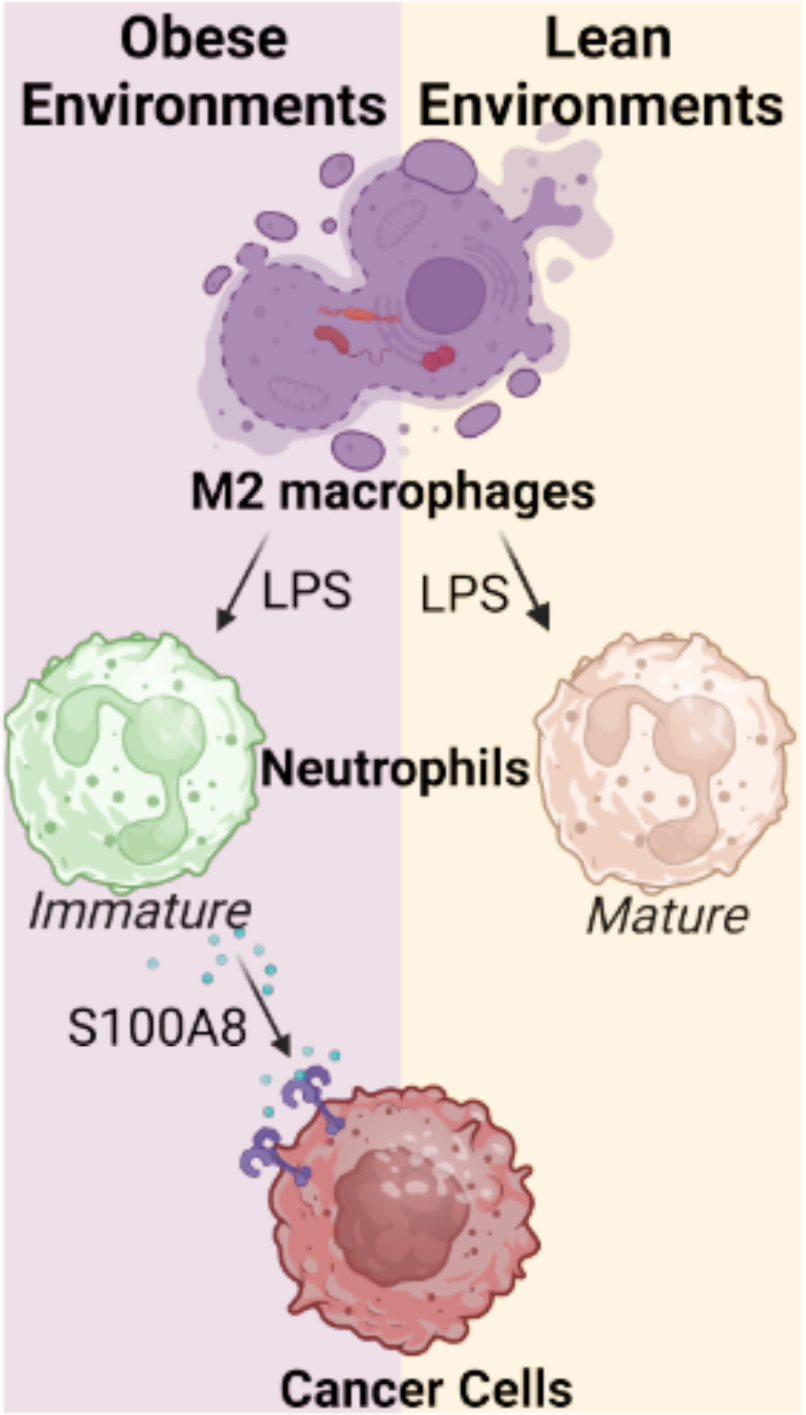
Model. A model of the cellular interaction network between M2-polarized macrophages containing gram-negative bacteria, immature neutrophils and cancer cells that drives obesity-induced post-menopausal breast cancer through TAN S100A8 release.

We highlight a role for GZMB positive iTANs in cancers and in particular obesity-induced breast cancers. While this cell population have gone largely unknown and uncharacterized, they have been reported in other unbiased IMC datasets using different GZMB antibodies for detection (Elaldi et al., 2021; Sheng et al., 2022), in orthoptic tumors after stimulation by lipid A – the hydrophobic group of LPS (Martin et al., 2018) and in granulomas from *Mycobacterium tuberculosis*-infected cynomolgus macaques (Mattila et al., 2015). Spatially, we demonstrate that GZMB^+^ TANs are restricted to areas close to dying macrophages that contain gram-negative bacteria supporting the requirement of LPS as stimulant for GZMB expression. Such niche-specific intrinsic flexibility of TANs aligns with recent work highlighting TAN reprogramming in hypoxic tumor regions leading production of secretory VEGF to enhance blood vessel formation (Ng et al., 2024). Similarly, our data places paracrine S100A8 signaling at the center of the interaction between TANs and cancer cells upon bacterial LPS stimulation in the obese environment. S100A8 is expressed abundantly by neutrophils and monocytes and is released during infection and inflammation through LPS, IFN-γ, IL-1β, and TNF-α signaling (Xu and Geczy, 2000; Xu et al., 2001). Interestingly, upon *in vivo* S100A8 blockade with neutralizing antibodies, LPS-induced neutrophil chemotaxis was prevented underscoring its potency on neutrophil biology (Vandal et al., 2003).

Neutrophil extracellular traps (NETs) are released in response to and help immobilize pathogens. With our experimental setup we were unable to consistently detect NET formation at the primary tumor site of either obese or non-obese breast cancer patients. This contrasts with what is found at the metastatic sites – especially the lung. The lung is particularly enriched in mature neutrophils (Sibille and Marchandise, 1993) that upon metastatic colonization is stimulated to release NETs in response to physiological stressors as obesity (McDowell et al., 2021; Quail et al., 2017) and stress (He et al., 2024).

Accumulating evidence suggest that bacteria are an integral part of the tumor microenvironment (Bullman, 2023). Consistent with an active role in the tumor, we demonstrate that higher tumor microbiome diversity is associated with improved survival in our hormone receptor negative breast cancer cohort. This aligns with previous findings in pancreatic cancer (Riquelme et al., 2019). To better understand what cell type in the tumors that harbor bacteria we resolved the LPS positivity across all identified cell clusters in our spatial analysis. This analysis revealed that M2-polarized macrophages contain the most LPS with 3.4% of the cells being positive. This is consistent with a recent spatial analysis of oral squamous cell carcinoma and colorectal cancer showing that the macrophages are the main cell type for *Fusobacterium* and *Treponema* positivity (Galeano Nino et al., 2022). Interestingly, despite being closely linked to leaky gut, our data suggest that the tumor microbiome is comparable between obese and non-obese breast cancer patients. The accumulation of LPS in tumor M2-macrophages is reminiscent of infection with *M. tuberculosis* – the causative agent of tuberculosis. *M. tuberculosis* infect host M2 macrophages in the lung to replicate in the phagosome (Bo et al., 2023). Here, virulent *M. tuberculosis* induces a necrotic cell death that fosters dissemination of newly formed bacteria whereas macrophages infected with attenuated strains of *M. tuberculosis* undergoes apoptosis and subsequent antigen presentation by dendritic cells (Behar et al., 2010). Our analysis cannot determine if the LPS positivity is based on macrophage engulfment of pathogens or active pathogen infection of macrophages. However, as the LPS positive M2-polarized macrophages in our analysis were also undergoing necrotic cell death this could suggest that gram negative bacterial infection of tumor associated macrophages is to some degree gaining fitness within the tumor ecosystem. At the same time, our data demonstrate that rather than having a direct beneficial effect on cancer cells, the LPS positivity endow the tumor a selective advantage through non-cancer cell autonomous regulation of S100A8 release by surrounding neutrophils. As such, our data supports a model of mutual beneficial relationship between prokaryotes and cancer cells within the tumor ecosystem.

In sum, our systemic analysis reveals that obesity-induced tumor growth is governed by a spatially restricted cellular network between LPS^+^ M2-polarized macrophages, iTANs, and cancer cells.

## Acknowledgements

We thank Erik Løkkevik, Bjørn Østenstad, Steinar Lundgren, Terje Risberg, and Ingvil Mjaaland for providing clinical samples. The mass cytometry was performed at the Flow Cytometry Core Facility, Department of Clinical Science, University of Bergen. Helios Mass Cytometer was funded by Bergen Research Foundation. We thank the genomic score facility (GSF) at the University of Bergen, which is a part of the NorSeq consortium, provided services on RNA-seq. GSF is supported by grants from the Research Council of Norway (245979/F50) and the Trond Mohn Foundation (BFS2016-genom). The immunofluorescent imaging was performed at the Molecular Imaging Center, Dept. of Biomedicine, University of Bergen. We thank Dr. Ann Richmond and Dr. Harold Moses for sharing the PyMT syngeneic cell line. We are grateful to the patients whose information and samples were vital for this study.

## Author contributions

Conceptualization: NH, STT

Methodology: STT, SMG, MEL, PL, WD, RKL, SK, PEL, JL

Funding acquisition: NH

Resources: NH, SK, PEL, JL

Supervision: NH

Writing – original draft: NH, STT

Writing – review & editing: NH, STT, JL, SK.

## Funding

The Norwegian Research Council, Young Talent #275250 (NH)

The Norwegian Research Council, Scientific Excellence #334437 (NH)

The Blix Foundation (NH)

The Meltzer Foundation (STT)

Centre for Molecular Medicine Norway, Seed Funding (NH, STT)

## Data and materials availability

The raw RNA-seq data for this manuscript are available through European Nucleotide Archive (ENA) at EMBL-EBI under accession number PRJEB75365.

All imaging mass cytometry images are available through Zenodo accession number 10.5281/zenodo.11072468.

R-script for cell mask conversion into REDSEA appropriate format and codes related to the adipocyte detection algorithm are available are available at GitHub (HalbergGroup).

## METHODS

### Clinical breast cancer samples

A total of 223 patients with primary stage III breast cancers were recruited during the trial inclusion period, between November 24^th^, 1997 and December 16^th^, 2003. Patients were randomized to neoadjuvant monotherapy with either epirubicin (90mg/m^2^) or paclitaxel (200mg/m^2^). All samples used in the present study were collected prior to treatment. All 37 post-menopausal (age >50 years) breast cancer patients included in this study were diagnosed with stage III primary breast cancer and where all estrogen receptor (ER) and progesterone receptor (PR) negative. Tumor size at diagnosis, gene expression by microarray at diagnosis, and survival rates were available for each patient. Approval of the clinical study was given by REK-Vest, Norway (Approval number 273/96-82.96). To generate tissue microarrays (TMAs) used for IMC and IF experiments, 5 mm sections of paraffin cores with a diameter of 1.2 mm were included per patient. More details can be found in (Chrisanthar et al., 2011). For the IMC experiment, we included 1-3 cores per patient.

### Animals

All animal experiments were approved by the Norwegian Food Safety Authority (FOTS number 14041 and 30403) and conducted according to the European Convention for the Protection of Vertebrates Used for Scientific Purposes, Norway. The C57BL/6J mice were bred at the animal facility at the University of Bergen. The experiment only included female mice that were housed together (5-6 per cage) in IVC-II cages (SealsafeÒ IVC Blue Line 1284L, Tecniplast, Buguggiate, Italy) using standard housing conditions at 21 °C ± 0.5 °C, 55% ± 5% humidity, and 12 h artificial light-dark cycle (150 lux). During breeding and before experiments they received standard rodent chow (Special Diet Services, RM1 801151, Scanbur BK, Oslo, Norway) and water ab libitum. At 6 weeks old, the mice were randomly assigned to either a high-fat diet (HFD; D12492, Research diets) or a Chow diet (Special Diet Services, RM1 801151, Scanbur BK, Oslo, Norway) for at least 10 weeks before implantation of cancer cells. For RNA-seq, Seahorse, and CO_2_ Trapping, mice were fed HFD or Chow diet for 18, 11, or 10 weeks respectively before implantation with PyMT. Mice implanted with EO771 cells were fed HFD or Chow for 12 weeks before implantation. Throughout the tumor experiment the mice continued to receive their respective diets.

### Cell lines

PyMT (Sai et al., 2017) and EO771 (CH3 BioSystems) cells were cultured in DMEM with high glucose (D5671-500mL, Sigma-Aldrich), supplemented with 10% FBS ((v/v), F7524, Sigma-Aldrich), 1% Penicillin-Streptomycin ((v/v) P0781-100ML, Sigma-Aldrich)), and 1% L-Glutamine solution ((v/v) G7513-100ML, Sigma-Aldrich) in 5% CO_2_ and 37°C.

### Mammary-fat pad injections

1000 PyMT or 10 000 EO771 cells were orthotopically injected into the fourth inguinal mammary fat pad (MFP) of obese and non-obese female C57BL/6J mice. In preparation for injections, cells were diluted in PBS and thereafter mixed 1:1 with Matrigel (356231, Corning). The total injection volume was 50 µl. Mouse body weight was measured at the day of injection.

### Mouse tissue collection

MFPs and livers from HFD and Chow fed mice were collected. The MFP weight was measured and analyzed using GraphPad Prism (v.10.1.2). Livers were fixed in 4% PFA in PBS for one day, subsequently washed 2 times in PBS before they were stored in 70% EtOH until paraffin embedding. At last, the livers were sectioned (5 µm), stained with hematoxylin and eosin, and imaged using Olympus VS120 S6 Slide scanner with a 40x magnification.

### Imaging mass cytometry

#### Antibody labeling

Antibody conjugation was performed using Maxpar X8 Antibody Labeling Kit (Fluidigm) and MIBItag Conjugation Kit (600157, IonPath) according to manufacturer’s protocol.

#### Tissue staining

10 formalin-fixed paraffin embedded human breast cancer TMAs were stained according to Fluidigm’s Imaging Mass Cytometry Staining protocol (PN 400322 A3) with some modifications. Briefly, the tissue slides were baked at 60°C for 2 hours. Thereafter, deparaffination and rehydration were performed by placing the tissue slides in 2x xylene for 10 min each, and decreasing concentration of ethanol (100%, 96%, 80%, and 70%) for 5 min each. Further, the tissue slides were placed in Milli-Q water for 5 min. Antigen retrieval was performed at 95-98°C for 30 min using 1X of Target Retrieval Solution pH9 (S236784-2, Agilent). The tissue slides were cooled down for 20 min at RT, before they were washed with Milli-Q water and then PBS for 10 min each. To allow for proper tissue staining and blocking the tissues were encircled with a ImmEdge Hydrophobic Barrier PAP Pen (H-4000, Vector Laboratories), before blocking the tissues with 3% BSA in PBS for 45 min. Antibodies were spun down at 13 000xg for 2 min prior to antibody cocktail preparation. The antibody cocktail was made by diluting the antibodies (Table S1 and S2) in 1% BSA in PBS. The tissue slides were incubated with the antibody cocktail in 4°C overnight. Further, the tissue slides were washed 2x in PBS-T (0.05% Tween 20 (v/v)) for 8 min each, and further in 2x in PBS for 8 min each. Subsequently, the tissue slides were stained with 0.3125 µM Cell-ID Intercalator-Ir (201192B, Fluidigm) in PBS for 20 min at RT. The tissue slides were washed with Milli-Q water for 5 min, and lastly air dried for 20 min at RT.

#### Data acquisition

Data acquisition was performed at the Flow Core Facility at the University of Bergen utilizing the Hyperion Imaging System (Fluidigm) and the CyTOF software (Fluidigm). The tissue cores were ablated with a pixel resolution equal to 1 µm^2^. The ablation frequency was set to 200 Hz and the ablation energies used were 2 dB and 0 dB for IMC panel 1 and 2, respectively. 1-3 TMA cores per patient were ablated. The ablation area for most tissue cores were 848 µm x 848 µm. For tissue cores with missing or poor tissue quality, the ablation area was adapted. One small ROI of liver tissue was also ablated for all 10 TMAs, except for TMA9 in IMC panel 2.

#### Pixel classification and cell segmentation

The pixel classification and cell segmentation of adipocytes and the other cells within the breast TME were done separately, due to the morphological differences between these cell types.

##### IMC panel 1

The pixel classification and cell segmentation for IMC panel 1 were performed using the IMC segmentation pipeline (v.2.1 (Zanotelli and Bodenmiller, 2021)), including the use of the imctools python package (v.2.1 (Zanotelli and Bodenmiller, 2021)) and CellProfiler plugins called ImcPluginsCP (v.4.2.1 (Zanotelli et al., 2020)). The pipeline was followed as described in the documentation by the developers (Zanotelli and Bodenmiller, 2021). The imctools python script was run in Jupyter notebook (v.6.2.0) through launching the program through the imctools environment in Anaconda navigator (v.2.5.0). Since one of the MCD-files was corrupted, both zipped MCD-files and txt-files of the raw data were used to extract the data. A comma separated csv-file of the panel was made displaying which markers were included in pixel classification. The default settings were utilized to prepare the images for pixel classification using the CellProfiler pipeline named “1_prepare_ilastik”.

One of the cores was scanned in two sessions, because the machine lost power during ablation. The two ROIs were merged by importing the txt-files into R studio, using R (v.4.1.1 (Team, 2023)). The width varied between the two regions, the second region being 5 pixels wider, and were partially overlapping. To merge the ROIs, the first ROI was padded with columns of 0s to match the width, and rows overlapping with the first ROI were removed from the second ROI. The ROIs were then merged and saved as a txt file for further processing. The txt-file was also inspected and validated using MCD-viewer (v.1.0.560.2, Fluidigm).

###### Adipocytes in IMC panel 1

For the pixel classification and cell segmentation of adipocytes in IMC panel 1, only ROIs containing adipocytes were utilized (in total 29 ROIs). For most ROIs, pixel classification of adipocytes in Ilastik (v.1.3.3post1 (Berg et al., 2019)) was performed using the marker expressions of Perilipin-1 (In113 and In115) and Barium to identify the adipocytes. Barium was included because it was useful to distinguish between the pixels covering the other cells within the ROI and the area inside the adipocytes. For certain ROIs where it was challenging to identify the entire adipocyte membrane, Collagen I, and Ir193 were included as markers for the pixel classification. The entire ROI was used for pixel classification because random cropping of the images could result in exclusion of adipocytes. Pixels were labeled as “inside the adipocytes”, membrane, or background. For certain ROIs, where it was challenging to identify the adipocyte membrane, the image of the ROI before ablation visualized in MCD-Viewer (v.1.0.560.2 Fluidigm), was used as guidance for pixel labeling. Further, probability images for each ROI, generated after pixel classification in Ilastik (v.1.3.3post (Berg et al., 2019)), were utilized for cell segmentation in CellProfiler (v.4.0.7 (Stirling et al., 2021)). The CellProfiler pipeline “2_segment_ilastik” which was a part of the IMC segmentation pipeline (Zanotelli and Bodenmiller, 2021) was used with some modifications. In the “IdentifyPrimaryObject” module, several settings were changed. The “typical diameter of objects” was adapted to each specific ROI and the min value ranged from 2 to 50, while the max value ranged from 120 to 300. The “discard objects outside the diameter range” was set to “Yes”. The thresholding method was changed to “Manual” and equal 0.45. The “Automatically calculate minimum allowed distance between local maxima” was set to “No”, and a specific “suppress local maxima” value was decided for each ROI which ranged from 13 to 20. For the “IdentifySecondaryObject” module, the “method to identify the secondary objects”, was changed to Distance-B, with a 10-pixel primary object expansion. For certain ROIs, an “EditObjectsManually” module was added after the “IdentifyPrimaryObject” module to remove objects which were not adipocyte masks.

###### Non-adipocyte cells

66 randomly 500 x 500 pixels cropped ROIs of the patient breast cancer tissue were used for the pixel classification and cell segmentation of most cells in IMC panel 1. Most markers (Perilipin-1 (In113 and In115), aSMA, H3K4me1, CD14, CD16, SOX9, Pan-Keratin, CD11b, CD31, H3K27me3, CD44, CD11c, FoxP3, CD4, E-cadherin, CD68, CK19, CD20, CD8, CD133, CK7, pCREB, CD45RA, GZMB, Ki-67, Collagen I, CD3, pERK, CC3, CD45RO, Pan-actin, Histone H3, Ir191, and Ir193) were included for pixel classification using the Ilastik software (v.1.3.3post1(Berg et al., 2019)). All features with a σ from 1.00 – 10.00 were selected for pixel classification. To train the random forest classifier, pixels within the cropped ROIs were labeled according to which class they belonged to, nuclei, cytoplasm, and membrane or background. An uncertainty filter within the Ilastik software, and uncertainty images generated by the software was used as guidance for determining which pixels or ROIs to label, respectively, for efficient training of the classifier. For cell mask generation, probability images of the entire ROIs were imported into the CellProfiler software (v.4.0.7 (Stirling et al., 2021)) and the CellProfiler pipeline “2_segment_ilastik” (Zanotelli and Bodenmiller, 2021) were applied with some modifications. For optimal nuclei recognition several parameters within the “IdentifyPrimaryObjects” module were adapted. The pixels for the “typical diameter of objects” were selected for each individual ROI, where the min value was either set to 2 or 7, and the max value was set to 25. “Discard objects touching the border of the image” was set to “No”. Further, manual thresholding with specific thresholding value for each ROI was used, which ranged from 0.45 to 0.67. “Automatically calculate minimum allowed distance between local maxima” was set to “No”, and a “suppress local maxima” value was set for each ROI and was set to either 6 or 7.

##### IMC panel 2

###### Adipocytes

The adipocyte pixel classification and cell segmentation were performed utilizing the Steinbock pipeline (version 0.16.1 (Windhager et al., 2023)) for IMC panel 2. Only breast cancer ROIs with adipocytes were included in the pixel classification and segmentation. For optimal pixel classification, the ROIs were divided into 3 groups: 1) ROIs with adipocytes with clear membranes, 2) adipocytes with uncomplete boundaries and 3) ROIs with small adipocytes. A pixel classifier was generated for each of the groups. Prior to pixel classification the images were subjected to pixel filtration with a threshold of 50 to remove hot pixels. Pixel classification was performed using Ilastik (version 1.3.3post1(Berg et al., 2019)) The adipocyte markers CD36 and a merged channel of the Perilipin-1 (In113 and In115) were included in the pixel classification. In addition, to prevent segmentation of CD36^+^endothelial cells and macrophages, CD31 and CD68, were included. ColI and Ir193 were also included because the staining pattern of ColI and the background contrast and the overview of the cell dense area of Ir193 demonstrated the outline of the adipocytes and were therefore useful for recognizing the adipocytes. The full-size ROI image was used for pixel classification because randomly cropped training images could potentially exclude adipocytes. All features from 1.00 – 10.00 were selected for pixel classification. Three classes were used for pixel classification: 1) Inside adipocyte, 2) Membrane and 3) Background. The different marker expressions included in the pixel classification together with the guidance of the “Before Ablation Image” in MCD Viewer (v1.0.560.6, Fluidigm) were used to label the pixels into the three different classes. Further, cell masks were generated using CellProfiler (4.2.6 (Stirling et al., 2021)). Probability images of each ROI were imported into the Steinbock generated CellProfiler pipeline where some settings were changed in the “IdentifyPrimaryObjects”module for each ROI. For the “typical diameter of objects” the min value ranged from 5 to 50, while the max value ranged from 120 to 400. The “discard objects outside the diameter range” was set to “Yes”. “Manual” was selected as thresholding method and was set to 0.45. The “Automatically calculate minimum allowed distance between local maxima” was set to “No”, and the “suppress local maxima” value was set to 13 or 30. In the “IdentifySecondaryObjects” module, “Distance - B” with a pixel expansion equal to 10 was selected to find the cell boundaries within the tissues. For ROIs where the adipocyte masks were over-segmented or background was recognized as cells, the “EditObjectsManually” module was added in the CellProfiler pipeline to merge the cell masks or to remove cell masks, respectively.

###### Non-adipocyte cells

Pixel classification and cell segmentation of most cells within the breast TME was done using the Steinbock pipeline (version 0.16.1 (Windhager et al., 2023)) for IMC panel 2. As a part of the Steinbock pipeline hot pixel filtration was done with a threshold set to 50. Further, using Steinbock, training images of the ROIs were cropped with a size of 500 x 500 pixels, and the seed was set equal to 123. The cropped training images were further imported into Ilastik (1.3.3post1 (Berg et al., 2019)) for pixel classification. All features from 1.00 – 10.00 were selected. The expression of aSMA, CD15, CD14, MPO, CD16, CD163, Pan-keratin, CD11b, CD31, CD44, CD80, CD4, E-cadherin, CD68, CK19, CD20, CD8, CD206, CK7, CD45RA, GZMB, Ki-67, ColI, CD3, CC3, CD45RO, Pan-actin, HisH3, Ir191, and Ir193 were selected as markers for the pixel classification. The pixels in the cropped ROIs were labeled and placed into three different classes 1) Nuclei 2) Membrane/cytoplasm, and 3) background class, based on the marker expressions. The uncertainty image generated within Ilastik was also used as guidance to label pixels that the random forest classifier was uncertain about for the next iteration. Further, the classifier was applied on the full-sized ROIs and probability images were generated for each ROI. Cell segmentation was performed by importing the probability images into CellProfiler (version 4.2.6 (Stirling et al., 2021)). The Steinbock CellProfiler pipeline for cell segmentation was used with some modifications in the “IdentifyPrimaryObjects” module. The pixels for the “typical diameter of objects” were selected for each ROI, where the min was set to 2 or 7, while the max value was set to 25. “Discard objects touching the border of the image” was set to “No”. Further, manual thresholding with specific value for each ROI was used, where the value ranged from 0.45 to 0.65. Further, “Automatically calculate minimum allowed distance between local maxima” was set to “No”, and a “suppress local maxima” value was set for each ROI which ranged from 6 to 8. For some ROIs the “EditObjectsManually” module was used to remove masks within the probability map that were not nuclei or cells but recognized as that.

#### Combination of cell masks of adipocytes and most cells: IMC panel 1 and IMC panel 2

The cell masks of adipocytes and other cells were imported into CellProfiler (v.4.0.7 or v.4.2.6 (Stirling et al., 2021)) as tiff files. To make the masks become objects within CellProfiler, the adipocyte mask and the other cell masks were assigned with the new names “Adipocyte_mask” and “Other_cells_mask”, respectively, in the “NamesAndTypes” module using the setting “Assign a name to Images matching rules”. For IMC panel 2, the filter used to recognize the specific images in the “NamesAndTypes” module had to be changed to the new image names, which was changed because the Steinbock pipeline was used to analyze the images from this panel. Further, the masks were combined using the “CombineObjects” module, and the overlapping objects were handled by selecting “Segment”. Thereafter, the merged mask was converted from an object to an image with a uint16 color format using the “ConvertObjectsToImage” module. At last, the merged cell mask was saved as a 16-bit integer tiff image using the “SaveImages” module.

### Single cell analysis

#### Cores excluded from the analysis

##### IMC panel 2

For IMC panel 2 only breast cancer samples were analyzed. ROIs with decreasingly lower signal within the core were removed to be sure that only cores with representative staining were used for analysis. Also, one core that was ablated in another MCD-file which only consisted of one line of ablated tissue was not included in the analysis. Hence, in total 70 ROIs were included in the analysis from 35 patients (obese *n* = 22, non-obese *n* = 13). For some cells in one ROI only Ir191 and Ir193 signal were present, and therefore these specific cells were removed from the downstream analysis.

##### Compensation of cell mask spillover

For IMC panel 1, cell masks generated by CellProfiler (v.4.0.7 (Stirling et al., 2021)) were processed in R (v.4.1.1 (Team, 2023)) to convert cell masks into an format appropriate for REDSEA (Bai et al., 2021). A custom MATLAB script then combined the processed files into a single segmentationParams.mat as required by REDSEA. Markers used to perform the compensation using REDSEA were CD4, CD11b, CD14, CD68, CD3, CD20, CD8, CD16, CD31, CD44, CD45RA, CD45RO, E-cadherin, and CK19. “elementSize” was set to 4, while “Cross style” was selected as “elementShape”. The output fcs-file “dataRedSeaScaleSizeFCS.fcs” which contained the REDSEA compensated marker expressions scaled by cell size, were used for further analysis.

##### FCS-file concatenating

For IMC panel 1, FCS-files generated after compensation of cell mask spillover were imported and concatenated in R (v.4.1.1 (Team, 2023)) using the flowCore package (v.2.6.0, (Ellis et al., 2021)). Cells with a cell size below 10 were removed from the data set. X- and Y-coordinates were determined by locating the segmented cells in each cell mask using a custom R-script.

##### Read in single cell data, cell masks and images

For IMC panel 2, single cell IMC data, cell masks and images generated using the Steinbock pipeline (Windhager et al., 2023), were imported into R studio (v. 2023.09.1+494) as describe in the extended version of (Windhager et al., 2023). Briefly, the single cell data were stored as a SpatialExperiment object utilizing the imcRtools package (v.1.8.0 (Windhager et al., 2023)). Further, patient metadata; including patient id, ROI name, and condition were added to the SpatialExperiment object. Further, the expression counts for each marker were arcsinh transformed with a cofactor of 1. Stacked images of every marker expression of each ROI and the cell masks for every ROI were loaded into R studio utilizing the cytomapper package (v.1.14.0 (Eling et al., 2020)).

##### Compensation of metal spillover

Compensation of metal spillover for IMC panel 2 was performed following the instruction given in the extended version of (Windhager et al., 2023). Briefly, txt-files used for compensation were downloaded from https://zenodo.org/records/7575859 (v.0.1.2 (Eling and Windhager, 2023; Windhager et al., 2021)), arcsinh transformed with a cofactor set to 5, and utilized to generate a spillover matrix which was applied on the single cell data from IMC panel 2 using the CATALYST (v.1.26.0 (Crowell et al., 2023)) R package.

##### Uniform Manifold Approximation and Projection (UMAP)

###### IMC panel 1

UMAP was used for dimensionality reduction utilizing the uwot package (v.0.1.10, (Melville, 2020)) in R (v.4.1.1 (Team, 2023)). For reproducibility a seed was set to 42. Markers included for UMAP analysis were CD3, CD4, CD8, CD11b, CD14, CD16, CD20, CD31, CD44, CD45RA, CD45RO, CD68, CD133, CK7, CK19, E-cadherin, FOXP3, Granzyme B, HLA-DR, Pan-Keratin, and Perilipin_sum (sum of In113 and In115). Random noise (0-0.1) was added to the marker expression values, to reduce picket fencing, before being arcsinh transformed with a cofactor of 1. UMAP parameters were set to n_threads = 7, n_sgd_threads = 7, and verbose = T. UMAP plots were made using the ggplot2 package (v.3.3.5 (Wickham, 2016)).

###### IMC panel 2

UMAP was applied on single cells from IMC panel 2, using the scater R package (1.30.1 (McCarthy et al., 2017)) as described in the extended version of (Windhager et al., 2023). A seed = 42 was used. Marker expression was arcsinh transformed with a cofactor of 1. Markers included for the analysis were Perilipin-1 (both In113 and In115), aSMA, CD15, Vimentin, CD14, MPO, CD16, CD163, Pan-Keratin, CD11b, CD31, CD80, CD4, CD36, E-cadherin, CD68, CK19, CD20, CD8, CD206, CK7, CD45RA, Granzyme B, ColI, CD3, and CD45RO. UMAP1 and UMAP2 were used for visualization utilizing the dittoSeq package (v.1.14.0 (Bunis et al., 2020)).

##### Clustering using PhenoGraph

###### IMC panel 1

For clustering, PhenoGraph in the Rphenograph package (v.0.99.1 (Chen, 2015)) was used. For reproducibility the seed was set to 42. The markers used for clustering were CD3, CD4, CD8, CD11b, CD14, CD16, CD20, CD31, CD44, CD45RA, CD45RO, CD68, CD133, CK7, CK19, E-cadherin, FOXP3, Granzyme B, HLA-DR, Pan-Keratin, and Perilipin_sum (sum of In113 and In115). Random noise (0-0.1) was added to the marker expression values, to reduce picket fencing, before being arcsinh transformed with a cofactor of 1. The k nearest neighbor was set to 30. Cell clusters were named according to which markers they expressed and their spatial localization within the ROIs. The mean marker expression for each cell cluster was visualized in a heatmap using the heatmaply package (v.1.3.0, (Galili et al., 2017)). Each column was scaled so that the maximum value was set to 1. This was done to be able to identify the marker expression by the different cell clusters, and followingly phenotype each cluster. The total cell count of each PhenoGraph cell cluster was visualized using the ggplot2 package (v.3.3.5 (Wickham, 2016)). Spatial plots with cells colored according to PhenoGraph cluster membership were made using the EBImage package (v. 4.36.0 (Pau et al., 2010)) in R (v.4.1.1 (Team, 2023)).

###### IMC panel 2

Prior to clustering, cells with an area less than 10 were removed. Clustering of single cells for IMC panel 2 was performed by following the workflow described in the extended version of (Windhager et al., 2023). Clustering was done using the Rphenograph (v.0.99.1 (Chen, 2015)) R package where k was set to 30 and 42 was used as seed. Markers that were used for clustering were Perilipin-1 (labeled with In113 and In115), aSMA, CD15, Vimentin, CD14, MPO, CD16, CD163, Pan-Keratin, CD11b, CD31, CD80, CD4, CD36, E-cadherin, CD68, CK19, CD20, CD8, CD206, CK7, CD45RA, Granzyme B, Collagen I, CD3, and CD45RO. PhenoGraph clusters were named according to which markers they expressed. This was determine using heatmap with the marker expressions for each cluster generated using the dittoHeatmap function in the dittoSeq R package (v.1.14.0 (Bunis et al., 2020)) as guidance. In addition, the spatial localization of the cell clusters was inspected by coloring the cell masks for each ROI according to which cell cluster they belonged to using the plotCells function in the cytomapper (v.1.14.0 (Eling et al., 2020)) R package. Cell clusters with similar marker expressions were merged by assigning them with the same cell cluster name.

##### Manual curation of cell clusters for IMC panel 1

###### General manual curation of cell clusters

Similar cell clusters were merged by assigning them with the same PhenoGraph number in R (v.4.1.1 (Team, 2023)). Similarities between cell clusters were determined by using hierarchical heatmap (heatmaply package (v.1.3.0, (Galili et al., 2017))) of the cell clusters and marker expressions together with the spatial localization of the cell clusters as guidance.

###### aSMA^+^ cells

To improve the clustering of aSMA^+^ cells in IMC panel 1, PhenoGraph using the Rphenograph package (v.0.99.1, (Chen, 2015)) was applied to all cells within the data set with an aSMA expression above 5. To make sure that the aSMA threshold was reasonable, the total cell count of the filtered cells from each cell cluster were visualized using the ggplot2 package (v.3.3.5, (Wickham, 2016)). Further, CK7, CK19, E-cadherin, Pan-Keratin, aSMA, CD31, Vimentin, and ColI were used to cluster the filtered cells with PhenoGraph. These markers were selected to be able to distinguish between aSMA^+^ cells that were also positive for epithelial markers, endothelial markers, and ColI. For reproducibility, the seed was set to 42, and the k-nearest neighbor was set to 50. The new PhenoGraph clusters generated were visualized in a hierarchical heatmap, using the heatmaply package (1.3.0, (Galili et al., 2017)). The hierarchical clustering within the heatmap, the mean expression of each marker for each cell cluster, and spatial localization of the cell clusters within the ROIs determined which of the newly generated cell clusters that were merged. This resulted in four aSMA^+^ cell clusters. At last, an additional threshold was applied on the aSMA^+^ filtered cells. This was based on the spatial localization of the cells within the new cell clusters, where filtered cells with a CD31 expression greater than 0.7 were placed into CL33 (aSMA^+^ endothelial cells). Filtered cells with an E-cadherin expression greater than 1.3 were put into CL31 (Myoepithelial cells), while filtered cells with a ColI expression greater than 10 were placed into CL36 (aSMA^+^ColI^+^).

###### CD8^+^ T cells

To separate GZMB-expressing CD8 T cells from CD8 T cells, in IMC panel 1, cells from the CD8 T cells clusters from the first PhenoGraph round were filtered out. Further, PhenoGraph using the Rphenograph package (v.0.99.1, (Chen, 2015)) were applied on the filtered cells using these markers; CD3, CD8, and GZMB. The seed was set to 42, and the k-nearest neighbor to 50. Further, the resulting PhenoGraph clusters were manually merged based on their GZMB-expression, using hierarchical clustering within heatmap (using the heatmaply package (v.1.3.0, (Galili et al., 2017)), and the cells spatial localization within the ROIs. This resulted in one GZMB^+^ CD8 T cells cluster (CL47), and one CD8^+^ T cell cluster (CL45).

###### Macrophages

From the first round of clustering for IMC panel 1, using PhenoGraph, the macrophage cluster was found to express CC3^+^. However, when inspecting these cells spatially, not all macrophages were CC3^+^. Therefore, to distinguish CC3^+^ macrophages and macrophages, only cells from the macrophage cluster found in the first round of PhenoGraph clustering of all cells were filtered out. Thereafter, PhenoGraph using the Rphenograph package (v.0.99.1 (Chen, 2015)), was applied on the filtered macrophages utilizing a seed set to 42, k-nearest neighbor equal to 50, and the following markers: CC3, CD68, CD16, and CD14. By utilizing the dendrogram of the hierarchical heatmap, made using the heatmaply package (v.1.3.0 (Galili et al., 2017)), the mean marker expressions of the cell clusters, and the cells spatial localization within the tissue, the new cell clusters were manually merged. This resulted in a macrophage cluster (CL84) and a CC3^+^ macrophage cluster (CL85).

###### FOXP3^+^ cells

FOXP3^+^ cells were identified in several PhenoGraph clusters after clustering of all cells in IMC panel 1. To generate a regulatory T cell cluster, all cells with a FOXP3-expression above 2 were filtered out and placed into a new cluster. To check if the FOXP3 threshold and cell filtration were reasonable, an overview of which cell clusters the filtered cells were extracted from was made and inspected.

###### Manual curation of cell clusters IMC panel 2

To improve the clustering by separating mutually exclusive markers, manual curation of cells was done. All thresholds used were inspected in scatterplots where the marker of interest was plotted against a mutually exclusive marker. To verify that the filtration of cells using the specific thresholds was reasonable a bar plot displaying the cell count of the cells filtered out from each cell cluster was inspected. In addition, the cell masks from the cell type in focus were overlaid on the marker expression of the marker of interest in all ROIs to observe if they overlapped. This was done using the plotPixels function in the cytomapper (v.1.14.0, (Eling et al., 2020)) R package. The manual curation of cells was performed using R (v.4.3.2 (Team, 2023)).

###### CD68^+^ cells

To improve the clustering of macrophages (CD68^+^ cells), cells with a CD68 signal above the 95^th^ percentile of CD68 expression, and a signal below the 95^th^ percentile of the respective markers, MPO, CK19, E-cadherin and CD31, were filtered and assigned to a new PhenoGraph cluster. This cell cluster was merged with the other macrophage clusters by labeling them with the same name.

###### CD8^+^ cells

To improve the clustering of CD8^+^ T cells, cells with a CD8 signal above the 95^th^ percentile and a signal below the 95^th^ percentile of the markers CD68 and MPO, were filtered and assigned to a new PhenoGraph cluster. This cell cluster was merged with the other CD8^+^ T cell clusters by labeling them with the same name.

###### CD4^+^ cells

To improve the clustering of CD4^+^ T cells, cells with a CD4 signal above the 95^th^ percentile and a signal below the 95^th^ percentile of the markers: CD8, CD68, and MPO, were filtered and assigned to a new PhenoGraph cluster. This cell cluster was merged with the other CD4^+^ T cell clusters by labeling them with the same name.

###### CD20^+^ cells

To improve the clustering of CD20^+^ B cells, cells with a CD20 signal above the 95^th^ percentile and a signal below the 95^th^ percentile of the respective markers: CD8, CD68, CD4, and MPO, were filtered and assigned to a new PhenoGraph cluster named B cells.

###### Adipocyte filtering

To improve adipocyte classification, e.g. adipocytes lacking Perilipin-1 and CD36 signal, cells with an area greater than the 99.5^th^ percentile of all cells (599.645 µm) were studied spatially and assigned to the adipocyte cell cluster.

###### Separation of Ki-67^+^ and Ki-67^−^ cancer cells

To separate Ki-67^+^ and Ki-67^−^ cancer cells, cells already assigned as cancer cells through PhenoGraph clustering, with a Ki-67 expression above the 80^th^ percentile of Ki-67 expression of all cells were filtered and assigned to a new PhenoGraph cluster named Ki67High cancer.

###### CD206^+^ or CD163^+^ macrophages

To separate CD206^+^ and CD163^+^ macrophages from other macrophages, cells from macrophages clusters already generated with PhenoGraph clustering with an expression above the 80^th^ percentile of CD206 of all cells or an expression above the 80^th^ percentile of CD163 of all cells were filtered and assigned to a new PhenoGraph cluster named CD206^+^ or CD163^+^ macrophages.

###### aSMA^+^ cells

Upon spatial inspection of the PhenoGraph generated cell clusters, several of the aSMA^+^ cells were inappropriately clustered. To improve clustering, cells from clusters: 6 (aSMA^+^Vim+ColI^+^), 12 (aSMA^+^Vim^+^ColI^+^), 17 (aSMA^+^ColI^+^), 28 (aSMA^+^E-cad^+^), 15 (Endothelial), 19 (ColI^+^CD45RA^+^Vim^+^), and 20 (ColI^+^CD45RA^+^Vim^+^), were filtered and re-clustered with the arcsinh transformed (cofactor = 1) value of the markers: CK7, CK19, E-cadherin, Pan-Keratin, aSMA, Vimentin, Collagen I, CD45RA, and CD31. Additionally, k was set to 50 and seed to 3908. New cell clusters with a similar marker expression were manually merged, based on the mean marker expression of each of the cell clusters as visualized using dittoHeatmap in the dittoSeq (v.1.14.0, (Bunis et al., 2020)) R package. This yielded 5 cell clusters named Endothelial, aSMA^+^ColI^+^ aSMA^+^ CD45RA^+^Vim^+^, and Myoepithelial. Further, these new cell clusters were placed into the original spatial experiment file together with the other cell clusters.

###### Heatmap (IMC panel 2)

Heatmap of marker expressions for each cell cluster for IMC panel 2 was generated following the workflow followed by the extended version of (Windhager et al., 2023). The mean arcsinh transformed (cofactor = 1) marker expression for each cell cluster were visualized and row scaled from 0-1 in the heatmap using the DittoHeatmap function in the dittoSeq (v.1.14.0, (Bunis et al., 2020)) R package.

###### Cell cluster abundance

For IMC panel 1, the cancer cell clusters were merged into one cancer cell cluster. Further, the proportion each cell cluster made up in each patient was calculated using the compositions R package (v.2.0.2, (van den Boogaart et al., 2021)). To test for significant differences in cell cluster abundance between non-obese (BMI ≤ 25) and obese (BMI >25) patients Kruskal-Wallis rank sum test was performed for each cell cluster using the stats R package (v.4.1.1, (Team, 2023)).

###### Region analysis

For IMC panel 1, region analysis was performed to find regions within the ROIs with similar cell cluster composition. All cancer clusters were merged into one cancer cluster prior to the analysis using R (v.4.1.1(Team, 2023)). CytoMAP (Stoltzfus et al., 2020) was used to do the region analysis. The data was organized into a CytoMAP format using R, where a csv file was generated for each cluster for each ROI. CytoMAP was run in MATLAB (v. 9.6.0.1072779 (R2019a)). The data was imported into CytoMAP using “Load Multiple Samples”. Further, neighborhoods were defined using default settings. Clusters were then clustered into regions using default settings, except the number of regions which was manually set to 5. All cell clusters and all breast cancer samples were used to generate the regions. The result was visualized by generating a region heatmap (**Figure S2A**). The proportion of cells within each region for each patient was calculated using the compositions (v.2.0.2 (van den Boogaart et al., 2021)) R package. Plotting and statistical testing with the Mann-Whitney U test was performed in GraphPad Prism (v.9.3.1).

To compare the cell cluster abundance within each region between non-obese and obese patients, patients with a cell count less than 100 within at least one of the regions were removed. The proportion of the different cell clusters within each region were calculated using the compositions (v. 2.0.2 (van den Boogaart et al., 2021)) R-package. Statistical differences between the proportion of cell clusters between obese and non-obese patients were tested using Welch Two Sample t-test in R (4.1.1 (Team, 2023)). Log2FC was calculated for the mean proportion of cells within each cell cluster within each region for obese patients versus non-obese patients. The cell cluster abundance differences between non-obese and obese patients within each region were visualized using the ggplot2 R package (v.3.3.5 (Wickham, 2016)). Significant cell abundances (*P* value < 0.05) within each region were highlighted. For cell clusters where there were less than two observations in both or either of the conditions (non-obese and obese), no statistical test was performed, and hence the cell cluster abundance is not presented in the bubble plot.

###### Interaction frequency analysis

For IMC panel 1, a SingleCellExperiment object was made for the single cell data including all cells present in the breast tissue ROIs. Prior to this, all cancer clusters were merged into one cancer cluster (CL2), resulting in a total of 18 cell clusters. Further, a spatial graph was generated using the buildSpatialGraph in the imcRtools R package (v.1.0.2, (Windhager et al., 2023)) using expansion and a distance threshold of 20. To find the interaction counts between each cell cluster for each patient, the countInteractions function in the imcRtools R package was used. The parameter “group_by” was set to PatientID, “label” to PhenoGraph cluster and “method” to classic. Further, the mean interaction counts for the obese and non-obese patient group were calculated using R (v.4.1.1 (Team, 2023)). Also, the log2FC of the obese mean interaction counts versus non-obese mean interaction counts were calculated in R. Mann-Whitney U test in R was used to test for significant difference between the mean count interactions between obese and non-obese patients for each cell cluster. The results were visualized in a bubble plot using the ggplot2 R package (v.3.3.5 (Wickham, 2016)). Interaction plots, displaying the interactions between cell clusters spatially, were generated using the plotSpatial function in the imcRtools R package.

###### Adipocyte area frequency plot

To compare the adipocyte cell area from non-obese and obese patients, cells from the adipocyte cluster (CL20) in IMC panel 1 with an area above 65 µm from patients that were confirmed to have adipocytes present in their ROI were filtered out using R (v.4.1.1 (Team, 2023)) and exported as csv-files. Adipocyte cell areas from non-obese and obese patients were compared and visualized using GraphPad Prism (v.9.3.1). Normality was tested using Shapiro-Wilk test, and a two-tailed Mann-Whitney U test was used to statistically compare the cell areas between the two conditions.

###### Relative frequency of TAN cell area

The cell size of all the TANs (CL27), in IMC panel 1, within ROIs from obese and from non-obese patients were compared. To test for significant differences a two-tailed Mann-Whitney U test was conducted. The frequency plot and the statistical testing were done in GraphPad Prism (v.10.1.2).

###### LPS analysis

To get an insight into which cell types were infected with gram-negative bacteria, LPS^+^ cells were quantified across all cell clusters. This was done using the IMC panel 2 data set. Prior to the analysis one patient ROI was removed because the LPS signal within this ROI was from hot pixels. To distinguish LPS^+^ and LPS^−^ cells, a threshold equal to the 99^th^ percentile of the exprs LPS signal (0.6631991, arcsinh transformed LPS signal with a cofactor =1) for all cells within all patients was utilized. To verify if the threshold was reasonable, it was inspected by plotting LPS vs various markers in scatter plots. All LPS^+^ cells were also spatially inspected, where LPS signal within ROIs was overlaid with the cell masks of LPS^+^ cells to see whether they overlapped or not. This was done using the “plotPixels” function in the cytomapper (v.14.0 (Eling et al., 2020)) R package. The percentage of cells with LPS signal above this threshold within each cell type for each patient were calculated in R (v.4.3.2 (Team, 2023)). A two-tailed Mann-Whitney U test was performed to test for significant difference in the percentage of LPS^+^ cells between obese and non-obese patients for each cell type using GraphPad Prism (v.10.1.2).

For spatial visualization of CD206^+^/CD163^+^ macrophage cell mask overlaps with LPS signal within each ROI, cells belonging to this cluster were filtered. Further, the plotPixels function in the cytomapper (v.14.0 (Eling et al., 2020)) R package was utilized, where this cell cluster was set as “object” and LPS as “colour_by”.

###### NETs quantification

To quantitatively assess the spatial concurrence of neutrophils undergoing NETosis within tissue samples, we utilized K-dimensional (KD) trees for the systematic organization of two cellular markers indicative of NETosis. Specifically, this analysis was predicated on the spatial coordinate of the centroids of cells exhibiting the uppermost 99^th^ percentile of expression for citrullinated histone 3 (H3Cit) and myeloperoxidase (MPO). These coordinates were algorithmically structured into KD trees for each tissue sample, leveraging the KD tree implementation in SciPy (v1.10.1 (Virtanen et al., 2020)). It is important to note that the centroid-based method of coordinate representation, while facilitating computational analysis, does not delineate the actual perimeters of cellular structures. Preliminary tests determined that a proximity threshold of 7 µm from a given centroid was sufficient to encapsulate potential cellular interactions, thereby resolving the limitations of centroid approximation. On this basis, an overlap event was defined as the occurrence of spatial overlap between cells expressing H3Cit and MPO within the 7 µm radius. Number of NETs in obese and non-obese samples were quantified and compared using a Fischer’s Exact Test (GraphPad Prism v.10.1.0).

###### Raw images

Raw marker intensity images were generated using MCD Viewer (v.1.0.560.2 and v1.0.560.6, Fluidigm). Markers were thresholded to remove background noise.

### Immunofluorescence

#### Tissue staining

Immunofluorescence (IF) staining was performed on FFPE human breast cancer TMAs. Briefly, the slides were baked at 60°C for 2 hours. Deparaffinization was performed by 10 x 2 min incubation with xylene, 3×2 min with 100% EtOH and 96% EtOH, 3 min with 80% EtOH, and 5 x 2 min with PBS with 0.05% Tween 20 (PBS-T). Antigen retrieval was done using 1x of Target Retrieval Solution pH9 (Agilent Dako, S236784-2) at 95-98°C for 30 min, and subsequently cooled down at RT for 20 min. Further, the tissue was washed 5 x 2 min in PBS-T. Tissue was encircled with ImmEdge Hydrophobic Barrier PAP pen (Vector Laboratories, H-4000), and further blocked with 10% goat serum (ThermoFisher, 50062Z) for 1 hour at RT. Further, the tissue was stained with anti-CD11b antibody (1:500, clone EPR1344, ab133357, Abcam) or anti-LPS (1:100, clone WN1 222-5, Hycult BioTech, 182-HM6011-20UG) diluted in 1% BSA in PBS overnight at 4°C.

Tissue was further washed with PBS-T 4 x 5min, and subsequently incubated with goat anti-rabbit Alexa Fluor 555 (1:500, Life Technologies, A21428) or goat anti-mouse Alexa Fluor 555 (1:400, ThermoFisher, A21422) for 1 hour at RT in the dark. The tissue was washed with PBS-T for 4 x 5 min, further blocked with 10% goat serum for slides subsequently stained with CD56 or CD68, and 3% BSA in PBS for slides stained with MPO for 1 hour at RT. Further, the tissues were incubated with either CD56 (1:100, clone 123A8, Novus Biologicals, NBP2-34279), CD68 (1:400, clone KP1, Invitrogen, 14-0688-80) or MPO (1:200, Polyclonal Goat IgG, Biotechne, AF3667-SP) diluted in 1% BSA in PBS overnight at 4°C in the dark.

Tissues were washed with PBS-T for 4 x 5 min, and subsequently incubated with goat anti-mouse Alexa Fluor 647 (1:1000, Life Technologies, A21238) for CD56 and CD68 stained slides, and Donkey anti-goat Alexa Fluor 647 (1:1000, Invitrogen, A-21447) for MPO stained slide for 1 hour at RT. Tissue slides were washed 4 x 5 min with PBS-T, and subsequently incubated with Alexa Fluor 488 anti-Granzyme B antibody (1:200, clone EPR20129-217, Abcam, ab225472) for 1 hour at RT in the dark. Thereafter, the tissue was washed 4 x 5 min with PBS-T, and then incubated with DAPI (1:1000, Sigma-Aldrich, D9542-10MG) for 10 min at RT. The tissue was washed 3 x 5 min with PBS and thereafter mounted with Prolong Diamond Antiface Mountant (ThermoFisher, P36961). The tissue samples were placed in the dark at RT for at least 1 day before imaging.

The slides were imaged with 40x magnification using Olympus VS120 S6 Slide scanner at Molecular Imaging Center (MIC) at the University of Bergen. Images were subsequently visualized using QuPath (v.0.2.3 (Bankhead et al., 2017)).

Similar protocol was followed for tissues stained with anti-Cleaved Gasdermin E, anti-CD68 and anti-Cleaved Caspase-3. However, Target Retrieval solution, Citrate pH 6.1 (Agilent Dako, S1699) was used for antigen retrieval. The first staining of the primary antibodies CD68 (1:400, clone KP1) and cGSDME (1:100, clone E8G4U, Cell Signaling Technology, 55879) was performed simultaneously. The staining of the secondary antibodies, goat anti-mouse Alexa Fluor 647 (1:1000) and goat-anti rabbit Alexa Fluor 555(1:500), the day after was also done at the same time. In addition, right after the PBS-T wash after the last secondary antibody incubation, the tissues were stained with Alexa Fluor 488 anti-Cleaved caspase-3 (1:20, clone 5A1E, Cell Signaling Technology, 67308) for 1 hour in the dark.

#### Mean LPS quantification and correlation to cell cluster abundances

IF images of human breast cancer TMA cores stained with an anti-LPS primary antibody and a Goat-anti-mouse Alexa Fluor 555 secondary antibody were analyzed using QuPath (v.0.2.3 (Bankhead et al., 2017)). Each core was annotated with a smaller circle than the core, to exclude staining artifacts present at the edges of the core. Further, erythrocytes were removed from the annotated area. Patient cores with poor tissue quality were excluded prior to the analysis. This resulted in analysis of 39 patient cores between 25 patients. To find the mean LPS signal for the annotated area for each core, “Compute intensity features” was used. Default settings were utilized, except that “Mean” was selected as “Basic features”. Further, the measurements of each annotation were exported from QuPath and imported into MS Excel. The mean LPS signal for each patient was calculated using MS Excel. The mean LPS signal for each patient was correlated with the IMC panel 1 cell abundance proportion for each patient where the IMC cluster abundance was known. The LPS stained cores and IMC panel 1 cores were from serial sections of the TMA. Spearman correlation was done using the cor and cor.test function in the stats R package (v.4.3.2 (Team, 2023)). The data was visualized in a bubble plot generated by using the ggplot2 (v.3.4.4 (Wickham, 2016)) R package.

#### Quantification of cGSDME^+^ macrophages within the breast TMEs

The number of CD68^+^CC3^+^cGSDME^+^ cells and CD68^+^CC3^+^cGSDME^−^ cells in patient TMA cores (*n* = 16 cores in *n* = 4 patients) stained for IF were manually counted using QuPath (v.0.5.0 (Bankhead et al., 2017)). The percentage of CD68^+^CC3^+^cGSDME^+^ cells out of all counted cells was visualized using GraphPad Prism (v.10.1.2).

#### Neutrophil assays

##### Tumor dissociation

Tumors from mouse mammary-fat pad (MFP) were harvested and placed in MACS Tissue Storage Buffer (Miltenyi Biotec, 130-100-008) on ice. Briefly, the tumors were dissociated by cutting the tissues into small pieces, and thereafter treated with RPMI 1640 (Sigma-Aldrich, R8758-500ML), enzyme A, D, and R as described in the Tumor Dissociation Kit manual (Miltenyi Biotec, 130-096-730). Further, the tumor was incubated in a rotisserie at 40 rpm for 30-40 min at 37°C. Thereafter, the cells were poured through a 70 µm strainer, and washed with 7.5 mL RPMI 1640 supplemented with 0.25 mg/mL DNase (Sigma-Aldrich, DN25-1G), and kept at RT for 10 min before being placed on ice. Then centrifuged at 300 g for 10 min. To remove red blood cells, the cells were incubated with 1x Red Blood Cell Lysis Solution (Miltenyi Biotech, 130-094-183) for 2 min at RT, and subsequently centrifuged at 300 g for 1 min. Lastly, the cells were resuspended in AutoMACS Rinsing Solution (Miltenyi Biotec, 130-091-222) supplemented with MACS BSA Stock Solution (Miltenyi Biotec, 130-091-376).

##### Neutrophil isolation

Neutrophils from dissociated tumors were isolated using the Neutrophil Isolation Kit (Miltenyi Biotec, 130-097-658), according to the manufacturer’s protocol.

##### Mitochondrial stress test - seahorse

To study the effect of S100A8 and S100A8/9 on murine breast cancer cells mitochondrial activity, 2× 10^4^ EO771 and PyMT cells were cultured in 100 µL of DMEM supplemented with 10% FBS and 4 mM glutamine with or without S100A8 or S100A8/9 recombinant protein (5 or 10 µg/mL) and plated in XFe96/XF Pro PDL Cell Culture Microplates (Agilent Technologies, Cat. #103799-100, US). Following 16h overnight incubation, the media was removed, washed with dulbecco’s phosphate buffered saline (DPBS: Merck, Cat. #D8662, Germany) twice, and 175 µL mitochondrial unbuffered, phenol-red free DMEM media (Merck, Cat. #D5030, Germany) supplemented with 10 mM glucose (Merck, Cat. #G8644, Germany), 2 mM sodium pyruvate (Thermo Fisher, Cat. #11360070, Norway), and 4 mM glutamine at pH 7.4 (+/−0.05) was added to each well. The plates were, then, transferred to a 37°C incubator not supplemented with CO_2_ for 1 h prior to mitochondrial stress assay. Throughout the mitostress experiments, 3 µM oligomycin (Merk, Cat. #75351, Germany), 1 µM Carbonyl cyanide 3-chlorophenylhydrazone (CCCP; Merck, Cat. #C2759, Germany), 1 µM rotenone (Merck, Cat. #557368, Germany), and 1 µM antimycin A (Merk, Cat. #A8674, Germany) were used.

To study the mitochondrial activity of neutrophils isolated from lean and obese mice, the protocol from (Monteith et al., 2022) was followed with a slight modification. Briefly, Corning Cell-Tak solution (final concentration is 22.4 µg/mL diluted in sterile Milli-Q water; Life Sciences, Cat. #354240, US) was prepared and 10 µL of the solution were applied to each well of X96 Seahorse plates. 96XF seahorse plates were, then, added 20 µL of sodium bicarbonate (NaHCO_3_, Merk, Cat. #S5761, Germany) buffer (pH 8.0) and incubated for 20 min at room temperature for solution adsorption. After incubation, the solution was aspirated from each well, washed with sterile Milli-Q water (200 µL/well) twice, and aspirated prior to the addition of neutrophils. 2×10^5^ neutrophils were resuspended in 50 µL neutrophil mitochondria assay media containing Seahorse Agilent RPMI media (Agilent Technologies, Cat. #103576-100, US), 1 mM pyruvate, 10 mM glucose, and 2 mM glutamine at pH 7.4 (+/−0.05) and added to each well. No neutrophils were added to the four wells in the corners of the plate as a background control. The plates were centrifuged at 200 x g (using zero braking setting) for 1 min. The plates were, then, transferred to a 37°C incubator not supplemented with CO_2_ for 30 min to ensure that the cells have completely attached. 130 µL of neutrophil mitochondria assay media was slowly added along the side of each well and the plates were returned to the incubator without CO_2_ for 15 min. For the mitochondrial stress test, the sensor cartridge (Agilent Technologies, Cat. #103775-100, US) was sequentially loaded with oligomycin (15 µM), CCCP (15 µM), rotenone (5 µM), and antimycin A (5 µM) (diluted in mitochondria assay media) for injection into the cell culture. Oxygen consumption and extracellular acidification rates were collected using a Seahorse XFe96 Analyser and analysed using Wave Desktop Software (Agilent Technologies, US). Protein measurement was performed for data normalization. The cells were washed twice with DPBS, lysed cells by freezing the plate at −80 freezer, and protein was measured using Pierce® BCA Protein Assay Kit (Thermo Fisher, Cat #23225, Norway). Plots were made in, and statistics were performed using GraphPad Prism (v.10.1.2).

##### CO_2_ Trapping

Glucose and palmitic acid (PA) substrate oxidation rate of neutrophils isolated from lean and obese mice was based on (Wensaas et al., 2007) and performed as in (Liu et al., 2022) with a slight modification. Briefly, CO_2_ trapping solution was prepared as follow: radiolabeled [1-^14^C] PA (2 µCi/mL; Perkin Elmer, Cat: #NET043001MC, US) and D-[^14^C(U)] glucose (2 µCi/mL; Perkin Elmer, Cat: #NEC042V250UC, US) were diluted in DPBS supplemented with 10 mM HEPES (Merck, Cat. #H0887, Germany) and 1 mM L-carnitine (Merk, Cat. #C0283, Germany). Respective amounts of non-radiolabeled substrate were added to obtain final concentrations of D-glucose (10 mM) and BSA-conjugated PA (200 µM). 96 well microplates (Thermo Fisher, Cat. #167008, Norway) were coated in Cell-Tak solution and sodium bicarbonate buffer, aspirated, washed with Milli-Q water twice, and aspirated prior to the addition of neutrophils. 2×10^5^ neutrophils were resuspended in 25 µL PBS supplemented with 10 mM HEPES and 1 mM L-carnitine and added to each well. The plates were centrifuged at 200 x g (using zero braking setting) for 1 min. 25 µL of CO_2_ trapping solution was added to each well to obtain final concentrations of radiolabeled [1-^14^C] PA (1 µCi/mL) and D-[^14^C(U)] glucose (1 µCi/mL) as well as respective amounts of non-radiolabeled substrate could obtain final concentrations of D-glucose (5 mM) and BSA-conjugated PA (100 µM). An UniFilter®-96w GF/C microplate (Perkin Elmer, Cat. #6055690, US) was activated for capturing the CO_2_ by the addition of 1 M NaOH (25 μL/well) and sealed to the top of the 96-well tissue culture plates and incubated for 5 h at 37°C. Subsequently, 30 µL scintillation liquid (MicroScint-PS, Cat. #6013631, PerkinElmer, US) was added to the filters and the filter plate was sealed with a TopSealA (PerkinElmer, Cat. #6050185, US). Radioactivity was measured using MicroBeta2 Microplate Counter (PerkinElmer, US). Protein measurement was performed for data normalisation. The cells were washed twice with DPBS, lysed by 0.1 M NaOH (Merck, Cat. #655104, Germany), and protein was measured using Pierce® BCA Protein Assay Kit (Thermo Fisher, Cat. #23225, Norway). Data were visualized and statistics were performed using GraphPad Prism (v.10.1.2).

##### Neutrophil culturing with LPS

Isolated neutrophils from tumors were cultured in a 96-well plate (100 000 cells/well) in 250 µl RPMI 1640 medium (Sigma-Aldrich, R8758-500ML) supplemented with 10% FBS ((v/v), F7524, Sigma-Aldrich) and 1% penicillin-streptomycin ((v/v), P0781-100ML, Sigma-Aldrich). To stimulate the neutrophils, 2 µl/mL LPS (*Escherichia coli* O26:B6, ThermoFisher, 00-4976-93) were added to the wells before incubation in 5% CO_2_ at 37°C for 18 hours.

##### RNA-extraction

Neutrophils were collected after 18 hours in culture and RNA was extracted using the Total RNA Purification Kits (Norgen Biotek, 37500) following the manufacturer’s protocol.

#### S100A8 cancer cell assays

##### Proliferation assay

To study the effect of S100A8 (R&D Systems, Cat. #9877-S8, US) and S100A8/9 (R&D Systems, Cat. #8916-S8, US) on breast cancer cell proliferation, 1×10^3^ EO771 and PyMT murine breast cancer cells were cultured in 100 µL of Dulbecco’s Modified Eagle’s Medium – high glucose (DMEM; Merck, Cat. #D5671, Germany) supplemented with 10% fetal bovine serum (FBS; Merck, Cat. #F7524, Germany) and 4 mM L-glutamine (Merck, Cat.G7513, Germany) with or without S100A8 or S100A8/9 recombinant protein (5 or 10 µg/mL) and plated in 96-well microplates (Thermo Fisher, Cat. #167008, Norway). Proliferation assay was conducted in The Sartorius IncuCyte S3 Live-Cell Analysis System (Ann Arbor, MI, US) at 37°C with 5% CO_2_ by imaging at 2h intervals using a 10x objective. The IncuCyte Zoom (v2022) software was used to calculate confluency values. Cell growth was determined by high content imaging and represented as ratio confluence normalized to the confluency value of the first image time point (reported as t=0). Data visualization and statistics were performed in GraphPad Prism (v.10.1.2).

##### Tumorsphere assay

To study the effect of S100A8 and S100A8/9 on tumorsphere formation, EO771 and PyMT cells were harvested using Accutase (Merck, Cat. #A6964, Germany) to obtain single cells and 1×10^4^ cells were re-suspended in 1 mL of standard stem cell media comprising DMEM F12 (Merck, Cat. #D8437, Germany) supplemented with mouse basic fibroblast growth factor (mbFGF, 20 ng/mL; Thermo Fisher, Cat. #PMG0034, Norway), mouse epidermal growth factor (mEGF, 20 ng/mL; Thermo Fisher, Cat. #PMG8044, Norway), and 1x B27 supplement (Thermo Fisher, Cat. #12587010, Norway). 1×10^4^ cells were diluted in 10-fold serial dilution to reach the number of 1×10^3^ in 1 mL stem cell media; 150 µl of the cell mixture (150 cells/well) were supplemented with or without S100A8 or S100A8/9 recombinant protein and plated to 24 corning corstar ultra-low attachment multiple well plates (Merck, Cat. #CLS3473, Germany). Tumorsphere were imaged using IncuCyte S3 Live-Cell Imaging Analysis System (Ann Arbor, MI, US) and counted after 5-8 days of growth. Graph generation and statistics were performed in GraphPad Prism (v.10.1.2).

#### mRNA extraction, cDNA synthesis, qPCR and primer design

Total RNA was isolated from approximately 600 000 neutrophil cells using the Qiagen total RNA extraction kit (Qiagen, Cat. #74106, US) following product guidelines, including on column digestion of gDNA. The concentration of isolated RNA was determined spectrophotometrically, and RNA purity was evaluated by 260/230 nm and 260/280 nm ratios using a nanodropTM 2000/2000c Spectrophotometers (Thermo Fisher, Cat. #ND-2000, Norway). cDNA was synthesized from 200 ng RNA using 50 μM oligo(dT)20 primer (Thermo Fisher, Cat. 18418020, Norway), 10 mM dNTP Mix (Thermo Fisher, Cat. 18427013, Norway), and SuperScript™ IV Reverse Transcriptase (Thermo Fisher, Cat. 18090010, Norway) as well as RNAaseOUT recombinant ribonuclease inhibitor (Thermo Fisher, Cat. 10777019, UK). Primers were designed using NCBI BLAST primer design or, where transcripts were not available in the NCBI database, primer 3 software was used (Untergasser et al., 2012). Amplicon size was restricted between 80 to 120 bp and amplicons were required to span an exon-exon boundary. The standard curve from 200 ng, 400 ng, 800 ng, and 1000 ng of each EO771 cDNA concentration ratio was carried to determine the efficiency of the primers. Primers were redesigned if amplification efficiency did not fall within 90-110% and/or a single peak was not observed in melting temperature analysis.

qPCR was carried out in two technical replicates with a LightCycler® 480 SYBR Green I Master Mix (Roche, Cat. #04887352001, US) using a LightCycler® 480 Instrument II (Roche, Cat. #05015243001, US). The results were calculated by ΔΔCt method using mouse actin (mActin) for mouse genes. Primer sequences are listed in Table S3. Plotting of data and statistics were done using GraphPad Prism (v.10.1.2).

#### RNA-sequencing and analysis

##### RNA sequencing

RNA extracted from PyMT tumor derived neutrophils from obese and lean mice were sequenced. Read alignment was done using HiSat2 v.2.0.5 (Reference genom: GRCm38). Samtools v.1.4.1 were used for indexing of bam. Reads were counted using subread-1.5.2. Reference transcriptome was gencode vM13q.

Principle Component Analysis (PCA) was performed on RNA-seq count values from untreated tumors derived HFD and Chow neutrophils using the prcomp function in the stats package (v.4.1.1). Prior to PCA, genes with a total count less than 5, across all samples, were removed and the counts were normalized and transformed using the rlog function in the DESeq2 package (v.1.34.0 (Love et al., 2014)). PCA was plotted using the autoplot function in the ggplot2 R package (v.3.3.5 (Wickham, 2016)).

The DESeq2 (v.1.34.0 (Love et al., 2014)) R package was used to test for differential expressed genes between the different tumor derived neutrophil conditions (HFD vs Chow) and treatment groups (untreated and LPS treated). Rows with a total gene count lower than 10 were removed before running DESeq2. Gene symbols were added to the data set using biomaRt (v.2.50.3 (Durinck et al., 2005; Durinck et al., 2009)) and the mmusculus gene ensemble dataset. All analyses were performed in Rstudio (v.2022.12.0) using R (v.4.1.1 (Team, 2023)).

#### Bubble plot of immature and mature genes

The log2FC values of HFD untreated vs Chow untreated genes calculated using DESeq2 (v.1.34.0 (Love et al., 2014)) of genes that have been reported to be related to immature and mature neutrophils, were visualized in a bubble plot. Only significant genes (adjusted *P* value ≤ 0.05) were included in the plot, and genes with an adjusted *P* value equal to NA were removed. For visualization, since the c-Kit adjusted *P* value was very small (adjusted *P* value = 0.000000e+00), the adjusted *P* value was set equal to the second smallest adjusted *P* value (adjusted *P* value = 3.869742e-101) which was the adjusted *P* value of Junb. The bubble plot was generated using the ggplot2 package (v.3.3.5 (Wickham, 2016)).

#### Gene Set Enrichment Analysis (GSEA)

Gene Set Enrichment Analysis (GSEA) (Castanza et al., 2023; Mootha et al., 2003; Subramanian et al., 2005) was performed on RNA-seq count values of untreated tumor derived neutrophils from HFD vs Chow fed mice. The count values were normalized prior to GSEA using the counts function in the DESeq2 (v.1.34.0 (Love et al., 2014)) R package. The normalized count expression data set in gct format and a phenotype label file in cls format were imported into GSEA (v.4.3.2). Default settings except the settings specified were used. The specific settings that were changed were as follows, “mh.all.v2022.1.Mm.symbols.gmt” was selected as “gene sets database”, the “permutation type” setting was set to “gene_set”, “Mouse_Ensembl_Gene_ID_MSigDB.v2022.1.Mm.chip” was selected as the “chip platform”, and “Metric for ranking genes” was set to “Diff_of_Classes”. The GSEA hallmark volcano plot was generated using the ggplot2 R package (v.3.3.5 (Wickham, 2016)).

#### Comparison of the effect of LPS on neutrophils isolated from obese and non-obese mice

Log2FC for HFD LPS vs HFD untreated, and Chow LPS vs Chow untreated were calculated using DESeq2 (v.1.34.0 (Love et al., 2014)). Further, genes with an adjusted *P* value of more than 0.05 were removed. To find the 99% prediction interval, a model was fitted and predicted by applying the lm and predict functions in the stat R package (v.4.1.1 (Team, 2023)), respectively, to the log2FC (HFD LPS vs HFD Untreated) and log2FC (Chow LPS vs Chow Untreated) values. Further, genes with a log2FC (HFD LPS vs HFD Untreated) and log2FC (Chow LPS vs Chow Untreated) above the upper 99% prediction interval were assigned as genes upregulated in the HFD condition. However, genes with a log2FC (HFD LPS vs HFD Untreated) and log2FC (Chow LPS vs Chow Untreated) below the lower 99% prediction interval were assigned as genes upregulated in the Chow condition. Genes with NA-values for log2FC (HFD LPS vs HFD untreated) or log2FC (Chow LPS vs Chow untreated) were excluded. The data was visualized using GraphPad Prism (v. 10.1.2).

#### GO: BP enrichment analysis

To investigate which pathways that the genes which were found to be up regulated in HFD LPS vs HFD untreated, compared to Chow LPS vs Chow untreated, a GO: BP enrichment analysis was performed. Briefly, genes above the 99% prediction interval were filtered out. The calculation of the prediction interval has been described earlier. The ENSEMBL gene name of these genes were found using the g: Convert function in gProfiler (Kolberg et al., 2023). To perform a GO: BP enrichment analysis on these genes, the enrichGO function in the clusterProfiler package (v 4.10.0 (Wu et al., 2021)) was used. The “organism” was set to “org.Mm.eg.db”, the “keyType” to “ENSEMBL”, “ont” to “BP” and the *P* value and the q value cut-off was equal to 0.05 and 0.10, respectively. The GO: BP enrichment dot plot was generated using the dotplot function in the enrichplot R package (v.1.22.0,(Yu, 2023)).

#### Secreted proteins

Genes which were found to have a log2FC (HFD LPS vs HFD untreated) and log2FC (Chow LPS vs Chow untreated) above or below the 99% prediction interval calculated, as described earlier, and that was either greater than 1 or less than −1 were filtered. DAVID (Huang et al., 2009; Sherman et al., 2022) was further used to determine which of these filtered genes that were containing a signal peptide and thus predicted to be secreted, which resulted in a total of 18 genes. Further, the mean TPM normalized RNA-seq counts of these genes for each sample group (HFD LPS, HFD, Chow LPS, and Chow) were calculated in R (v.4.3.2(Team, 2023)). A heatmap displaying the mean TPM normalized counts for each gene for each sample group was generated using the heatmaply R package (v.1.5.0 (Galili et al., 2017)), where values were scaled for each gene across each sample group.

#### 16S sequencing

The patient samples used in the present work were samples from postmenopausal patients with ER-negative, HER2-negative breast cancers. We included samples from all tumors (n=49) with these molecular characteristics drawn from a prospective trial (EpiTax) specifically designed for identification of predictive biomarkers for chemotherapy effect in breast cancer (Chrisanthar et al., 2011; Knappskog et al., 2012). For two patients (205 and 215) out of the 49, sufficient material for 16S sequencing analysis was not available, thus, the 16S data set included 47 patients.

##### Sample collection

Prior to commencement of chemotherapy, each patient in the EpiTax-trial had an incisional biopsy. Tumor tissue was snap-frozen in liquid nitrogen immediately upon removal in the operating theatre. Tissue was stored on liquid nitrogen until the time of further processing (DNA isolation).

##### Bacterial DNA extraction

Bacterial DNA was isolated from fresh frozen breast tissues (10-15 mg) using ZymoBIOMICS DNA Miniprep kit with the protocol specific for tissue samples and according to the manufacturer’s instructions (Zymo Research, Cat: #D4300). To minimize contamination, all pipettes, tips, and non-enzymatic kit components were UV-radiated for at least 1 hour prior to use, and extraction was performed in a dedicated pre-cleaned and irradiated laminar flow hood. The samples were digested for 60 minutes with proteinase K (10 µl) followed by bead-beating for 40 minutes (Vortex-Genie 2 with vortex adapter, Qiagen, Cat: #13000-V1-24) at maximal speed. DNA concentrations were measured by Qubit 2.0 fluorometer (Life Technology) using Qubit dsDNA HS Assay Kit (Invitrogen, Cat: #Q32584). The samples were enriched for microbial DNA using the NEBNext Microbiome Enrichment Kit (New England BioLabs, Cat: #E2612L), removing the methylated host DNA. Negative controls (nuclease-free H_2_O) were processed throughout the entire extraction procedure and included in the following 16S rRNA library preparations.

##### Library preparation and 16S rRNA gene sequencing

Amplification of targets for library preparation was conducted in two PCR steps. The first step was performed with primers targeting the V1-V2 region of the 16S rRNA gene (5’-TCGTCGGCAGCGTCAGATGTGTATAAGAGACAGAGMGTTYGATYMTGGCTCAG-3’ and 5’-GTCTCGTGGGCTCGGAGATGTGTATAAGAGACAGGCTGCCTCCCGTAGGAGT-3’). The V1-V2 primer set was chosen over the previously commonly used V3-V4 primer set, due to a considerably lower off target amplification of human DNA when sequencing the microbiota of human breast tissues (Walker et al., 2020). Pipettes and plasticware were UV-radiated before use, and PCR reactions were set up in a dedicated laminar flow hood. For PCR quality controls, TE-buffer was substituted for the DNA template. Amplification was performed in 25 µl reactions, containing 6.3 µl (25-100 ng) DNA template, 12.5 µl 1x NEBNext High Fidelity 2x PCR Master Mix (New England Biolabs, Cat: # M0541S) and 0.5M of each primer. Thermal cycling conditions were set as 98°C x 30 sec, followed by 28 cycles of 98°C × 10 sec, 62 °C x 30 sec, and 72 °C x 30 sec, then 5 minutes at 72°C. The PCR-products were purified with 20 µl AMPure XP beads (Beckman Coulter Genomics, Cat: #A63881) before performing the second PCR, with 8 cycles and using Nextera XT indexes (Illumina, Cat: # FC-131-1001) according to the 16S Metagenomic Sequencing Library Preparation Protocol from Illumina. After an AMPure XP bead cleanup, the libraries were validated with Bioanalyzer DNA 1000 (Agilent Technology), and concentrations were quantified using Qubit dsDNA HS Assay Kit. The libraries were pooled to equimolar ratios and diluted to 10 pM together with 15% PhiX and sequenced on the Illumina MiSeq platform using the V3 reagent kit (2 x 300 bp paired end).

##### Microbiome data analysis

16S rRNA gene V1-V2 libraries from 47 samples included in the present study and 35 samples from the same trial, not included in this study, along with 7 water-based negative controls and 7 other positive/negative controls, were sequenced in a single MiSeq run and analyzed together. Standard microbiome data analysis procedures were followed as outlined below:

1. Demultiplexed fastq files were initially trimmed of the V1-V2 primer sequences using cutadapt (Martin, 2011).
2. Forward and reverse reads were then filtered and trimmed of low-quality end bases using the dada2 R package (Callahan et al., 2016).
3. Dada2 was also utilized to construct PCR/sequencing substitution error models from all 82 samples, infer real biological amplicon sequence variants (ASVs), merge paired reads, and remove chimeras.
4. The Silva 138.1 prokaryotic SSU database (Quast et al., 2012) was employed to classify each unique ASV into taxonomy and assign (exact matching) species names.
5. ASVs unlikely to be contaminants from experimental procedures were identified based on the statistics of ASV prevalence in the 82 samples versus that in the 7 water-based negative controls using the decontam R package (Davis et al., 2018), with a default *P* value cutoff of 0.5.
6. Additional high stringency filters per run batch was set as ratio of reads in water-based negative controls versus reads in sample <0.5.
7. Alpha diversity (Shannon diversity) was determined using the R package Phyloseq (McMurdie and Holmes, 2013)

#### Survival analysis

##### S100A8 expression

Survival analysis of microarray S100A8 expression of human postmenopausal (≥ 50 years) hormone receptor negative (ER^−^PR^−^) breast cancer patients were conducted. The patients were separated based on BMI and their tumor S100A8 expression level. Patients with a BMI ≤ 25 kg/m^2^ were classified as non-obese, while patients with a BMI > 25 kg/m^2^ as obese. Further, a threshold equal to the median S100A8 expression (17.572) of all postmenopausal ER^−^PR^−^ patients (in total 44 patients) were utilized to separate S100A8^high^ (S100A8-expression > 17.572) and S100A8^low^ (S100A8-expression ≤ 17.572) expressing patients. The survival time in months and the Disease-Specific Survival (DSS) for each patient were used to generate the Kaplan-Meier plot. To test for statistical differences Log-rank Mantel-Cox was utilized. Hazard Ratio (logrank) of S100A8^high^ obese patients vs S100A8^low^ obese patients. Both plot generation and statistics were performed using GraphPad Prism (v.10.1.2).

##### Csn1s1 expression

Survival analysis of microarray Csn1s1 expression of postmenopausal (> 50 years) ER^−^PR^−^ breast cancer patients were performed. Patients were classified as obese (BMI > 25 kg/m^2^) or non-obese (BMI ≤ 25 kg/m^2^). Further, the patients with a Csn1s1 above the median (13.706) were classified as Csn1s1^high^, while patients equal to or below the median as Csn1s1^low^. Kaplan-Meier plots were generated using the DSS and survival time (months). Log-rank Mantel Cox was utilized to test for statistical significance. Hazard ratio (logrank) was calculated for obese Csn1s1^high^ vs obese Csn1s1^low^, and non-obese Csn1s1^high^ vs non-obese Csn1s1^low^. The statistics and graph generation were done utilizing GraphPad Prism (v.10.1.2).

##### Bacteria diversity

Survival analysis of bacteria genera diversity present within postmenopausal (> 50 years) ER^−^PR^−^ breast cancer patients were done. Patients were stratified only by the bacteria genera diversity or both the bacteria genera and BMI. Patients with an BMI > 25 kg/m^2^ were classifies as obese, while patients with an BMI ≤ 25 kg/m^2^ as non-obese. Patients were stratified based on the median Shannon diversity score (= 3.3128) of all patients. Patients with a diversity above or equal to median were classified as high diversity patients, while patients with a diversity below median as low diversity patient. Kaplan-Meier plots were made using relapse-free survival (RFS). Patients with no relapse free interval (*n* = 4) were not included. To test for statistical significance Log-rank Mantel Cox was used. Hazard ratio (log rank) was calculated for low diversity vs high diversity, The statistics were done, and graph made using GraphPad Prism (v.10.1.2).

#### METABRIC data

A dataset was constructed through the integration of survival data, BMI, and normalized gene expression levels and clinical information from data in the METABRIC cohort (Nguyen et al., 2023), resulting in the identification of 417 patients. Of these, 52 patients were post-menopausal, ER^−^ and PR^−^.

Survival outcomes and tumor size at the time of diagnosis within the patient cohorts stratified by either BMI (≥ 25) and the median expression level of S100A8. Kaplan-Meier survival plots were generated to visually represent the overall survival trajectories. Student t-tests were used to determine significance in tumor size. Cox proportional hazards regression was used to evaluate the mortality risk contingent on BMI and S100A8 expression.

##### Correlation calculation of S100A8 in METABRIC patients with known BMI

To determine the correlation between S100A8 gene expression and the expression of other genes within the METABRIC dataset we calculated Pearson correlation coefficients (S100A8 was excluded from the correlation analysis to avoid autocorrelation effects) and corresponding *P* values for each gene in relation to S100A8 expression. The results were organized and ranked by their correlation coefficients, enabling identification of genes significantly correlated with S100A8. This highlights genes with the most substantial correlation to S100A8, offering insights into their potential involvement in metabolic pathways or disease states associated with obesity.

#### ClusterProfiler GO: BP analysis of S100A8 correlated genes

Genes that were found to correlate more than R = 0.4 with S100A8 (total of 153 genes) in obese PM/ER^−^/PR^−^ from the Metabric patient data set were used to perform GO enrichment analysis. ENSEMBL gene names were found using the g: profiler g: Convert function (Kolberg et al., 2023), where “Human” was selected as organism. GO enrichment analysis was performed using the enrichGO function in the clusterProfiler (v.4.10.0 (Wu et al., 2021)) R package. Within the function, “org.Hs.eg.db” was set as “organism”, “keyType” was set to “ENSEMBL”, “ont” to “BP”, *P* value cut off to 0.05 and the q value cut off was set to 0.10. A dotplot was generated to visualize the results using the dotplot function in the enrichplot package (v.1.22.0 (Yu, 2023)).

#### Adipocyte Prediction using mask R-CNN

A dataset comprising 26 images from the IMC panel 1, each sized at 848 x 848 pixels (1 pixel = 1 µm^2^), was utilized for training the adipocyte detection model. These images were extracted from IMC data obtained from various patient states, including obese and lean conditions. The images were generated in R (v.4.3.1 (Team, 2023)) using the cytomapper package (v1.12.0 (Eling et al., 2020)) and annotated using a local client of the Computer Vision Annotation Tool (CVAT, v.2.7.0 (Sekachev et al., 2020)) to provide ground truth annotations of adipocytes. Ground truth was created by tracing the contours of the Perilipin 1 signal to provide approximate adipocyte masks. Annotations were exported in the Common Objects in Context (COCO 1.0) (Lin et al., 2014) annotation format.

The object detection model was trained using the Detectron2(v.0.6 (Wu et al., 2019)) in a python environment (v.3.8.17). Specifically, the model variant chosen was the mask region-based convolutional neural network (mask R-CNN (He et al., 2020), employing the ResNeXt (X101-FPN) architecture sourced from the Detectron2 model zoo API. The training duration was set to 300 000 epochs and was performed on a local machine with a NVIDIA GeForce RTX 4070Ti GPU.

During the training phase, a learning rate of 0.00025 was set, enabling a gradual convergence of the model. A learning rate decay strategy was implemented, halving the learning rate at specific epochs (80 000 and 180 000), to enhance the model’s adaptability and precision over the training duration.

The implementation of the adipocyte detection model was executed within the Python-based Detectron2 framework (v.0.6), which also facilitated functions for visualizing predicted adipocyte masks on processed images. Post-analysis involved the measurement and analysis of adipocyte mask sizes using scikit-image’s (v.0.19.3 (van der Walt et al., 2014)) regionprops and label functions.

#### Performance Evaluation of Adipocyte Prediction

A dataset comprising 15 images, each sized at 848 x 848 pixels, created for the purpose of evaluating the adipocyte detection model. These images were extracted from imaging mass cytometry (IMC) data obtained from various patient states, including obese and lean conditions. The images were generated in R (v.4.3.1 (Team, 2023)) using the cytomapper package (v1.12.0 (Eling et al., 2020)) and annotated using a local client of the Computer Vision Annotation Tool (CVAT) (Sekachev et al., 2020) to provide ground truth annotations of adipocytes.

The model’s performance was assessed using various standard evaluation metrics for image segmentation tasks such as F1 score and intersection over union (IoU) of the predicted adipocyte masks and the ground truth in the range of 0.50 to 0.95.

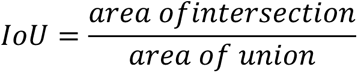

Evaluation was conducted on a dataset comprised of 15 annotated images with ground truth annotations of adipocytes.

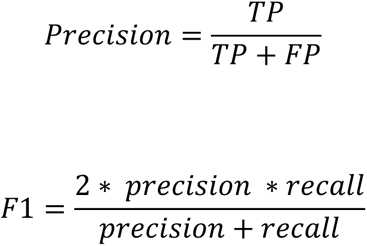

The model’s generalizability and ability to detect adipocytes was validated across two datasets obtained from different multiplexed imaging modalities: Imaging Mass Cytometry (IMC) (Jackson et al., 2020) and Multiplexed Ion Beam Imaging (MIBI) (Keren et al., 2018). This evaluation aimed to gauge the model’s ability in detecting adipocytes across diverse imaging technologies.

#### Adipocyte Classification in Validation Cohorts

Upon successful detection of adipocytes in validation cohort images derived from Multiplexed Ion Beam Imaging (MIBI) and Imaging Mass Cytometry (IMC), the size of each identified adipocyte was quantified for every image. In images containing three or more adipocytes and where the patients matched our patient cohort parameters (post-menopausal patients with triple-negative breast cancer), a median-based classification approach was employed to categorize the images being in either of the two size-based groups: “Small Adipocyte” and “Large Adipocyte”. Images exhibiting an average adipocyte size below the median were classified under “Small Adipocyte”, whereas those with an average size above the median were designated as “Large Adipocyte”.

#### Statistics

Please find specific statical test included in each method description. All statistics were performed in either R or GraphPad Prism. *P* < 0.05 were considered significant.

