## Supplemental Figures for "Interactions between neutrophils and macrophages harboring gram-negative bacteria promote obesity-associated breast cancer"

Takle *et al.*

This PDF include

Figures S1-S6

Tables S1-S3

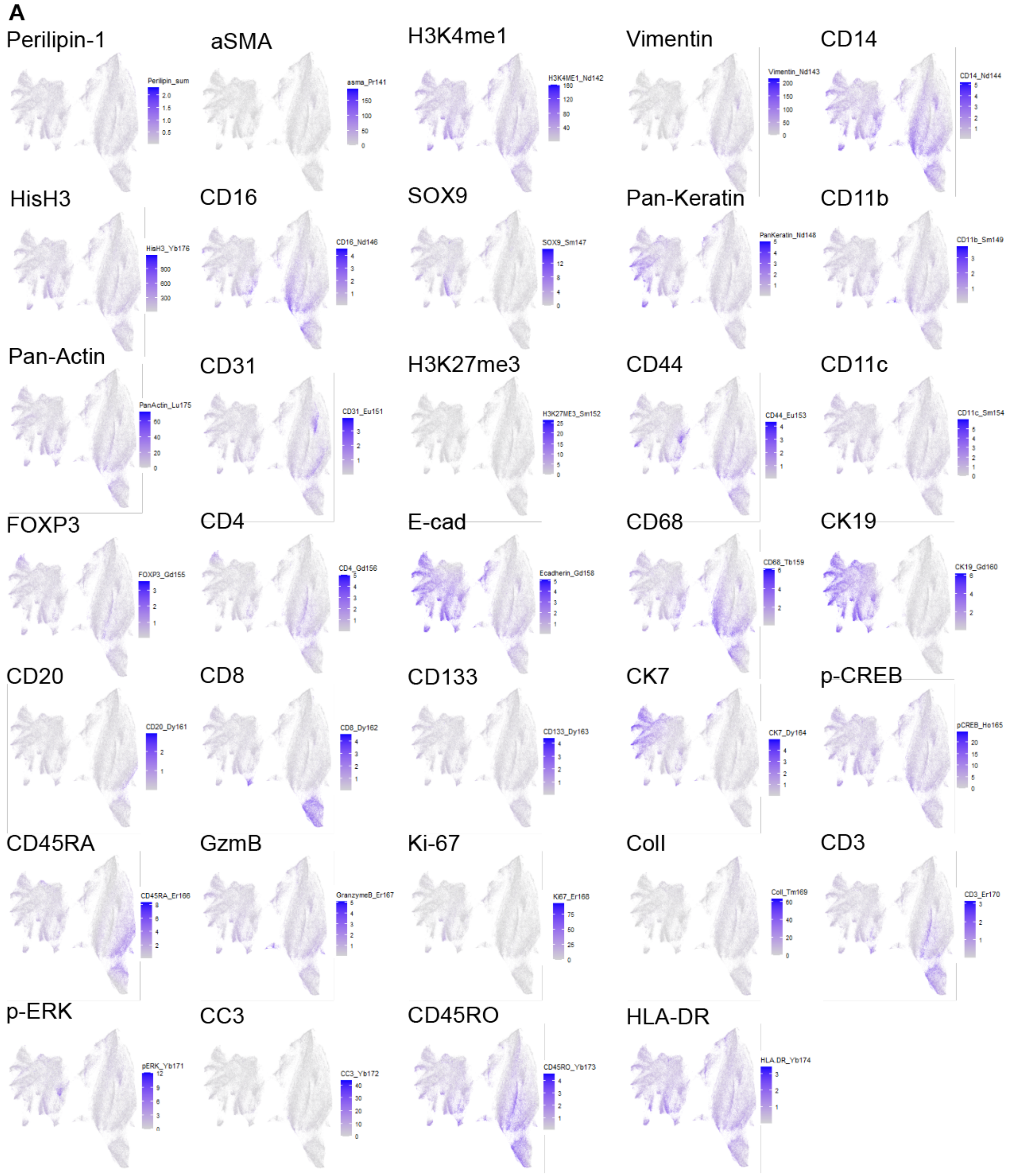
A
Perilipin-1
aSMA
H3K4me1
Vimentin
CD14
HisH3
CD16
SOX9
Pan-Keratin
CD11b
Pan-Actin
CD31
H3K27me3
CD44
CD11c
FOXP3
CD4
E-cad
CD68
CK19
CD20
CD8
CD133
CK7
p-CREB
CD45RA
GzmB
Ki-67
ColI
CD3
p-ERK
CC3
CD45RO
HLA-DR

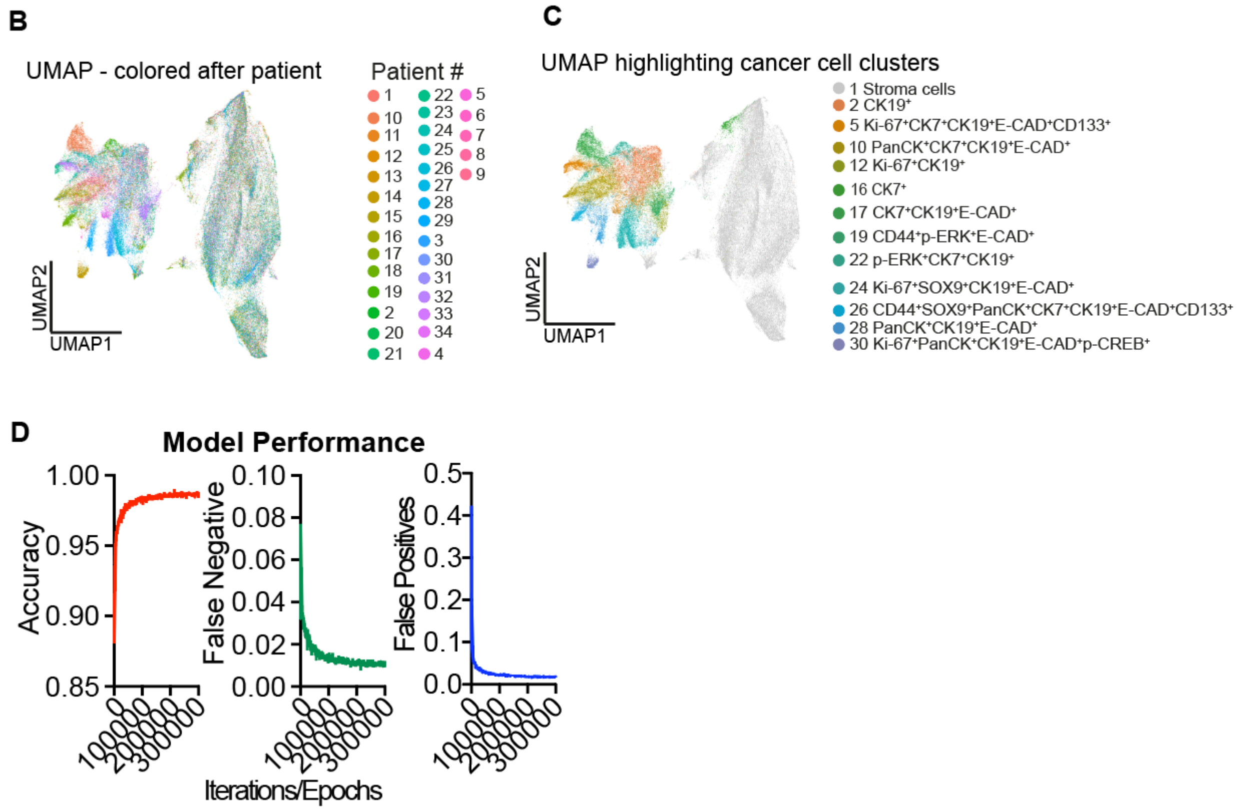


**Figure S1. Systemic characterization of the tumor microenvironment in post-menopausal and hormone receptor negative breast cancer.**

UMAP of cells in IMC panel 1 from obese (*n* = 22) and non-obese (*n* = 12) postmenopausal $ER^-$ $PR^-$ breast cancer patients colored by (**A**) marker expression, (**B**) Patient ID or (**C**) cancer clusters or stroma highlighting the different cancer cell clusters identified. (**D**) Training metrics of the computer vision-based adipocyte detection model during the training period of 300 000 epochs.

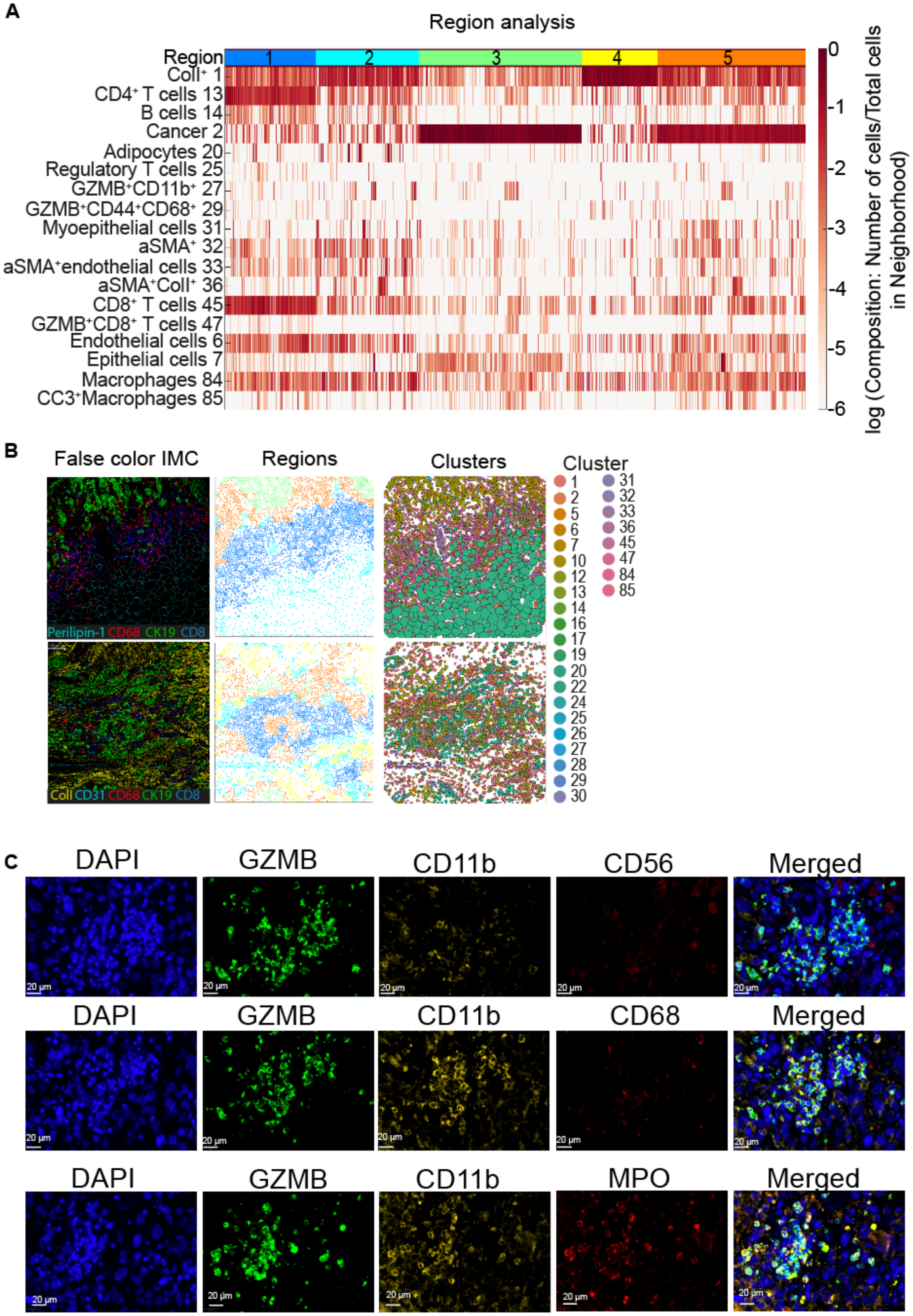
A
Region analysis
Region
1
2
3
4
5
ColI⁺ 1
CD4⁺ T cells 13
B cells 14
Cancer 2
Adipocytes 20
Regulatory T cells 25
GZMB⁺CD11b⁺ 27
GZMB⁺CD44⁺CD68⁺ 29
Myoepithelial cells 31
aSMA⁺ 32
aSMA⁺endothelial cells 33
aSMA⁺ColI⁺ 36
CD8⁺ T cells 45
GZMB⁺CD8⁺ T cells 47
Endothelial cells 6
Epithelial cells 7
Macrophages 84
CC3⁺Macrophages 85
0
-1
-2
-3
-4
-5
-6
log (Composition: Number of cells/Total cells in Neighborhood)
B
False color IMC
Regions
Clusters
Cluster
1 2 5 6 7 10 12 13 14 16 17 19 20 22 24 25 26 27 28 29 30
31 32 33 36 45 47 84 85
Perilipin-1 CD68 CK19 CD8
ColI CD31 CD68 CK19 CD8
C
DAPI
GZMB
CD11b
CD56
Merged
DAPI
GZMB
CD11b
CD68
Merged
DAPI
GZMB
CD11b
MPO
Merged
20 µm

**Figure S2.**

(**A**) Region analysis of cells in IMC panel 1. Heatmap of cell cluster composition within each region (1-5). Each line is colored from white to red by fraction each cell type makes up of all cells in each neighborhood. (**B**) Representative images of false color IMC marker expressions (Perilipin-1/CD31 = cyan, CD68 = red, CK19 = green, and CD8 = blue, ColI = yellow), the spatial localization of the regions (Region 1 = blue, Region 2 = cyan, Region 3 = Green, Region 4 = yellow, and Region 5 = orange) within the ROI and the spatial localization of the cell clusters within the ROI. (**C**) $GZMB^{+}CD11b^{+}$ cells within the breast tumor microenvironment are $MPO^{+}$. Serial sections of human breast cancer TMAs were stained with antibodies targeting GZMB (green), CD11b (yellow), CD56 (red), CD68 (red), and MPO (red), and stained with DAPI (blue) for IF.

A

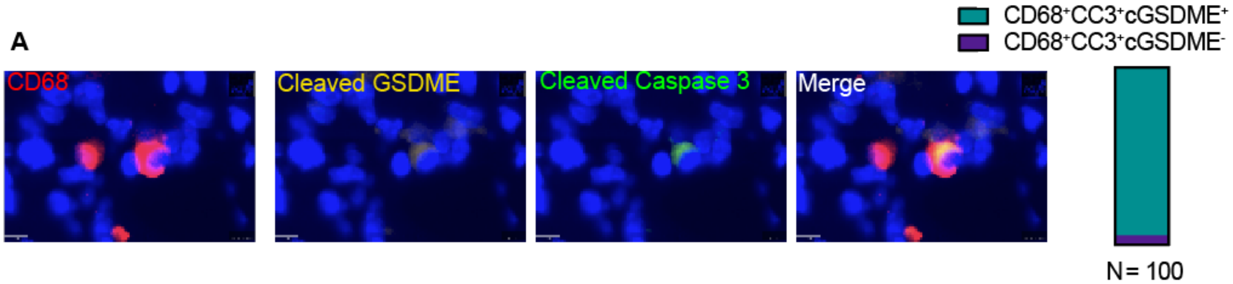
CD68
Cleaved GSDME
Cleaved Caspase 3
Merge
CD68+CC3+cGSDME+
CD68+CC3+cGSDME-
N= 100

B

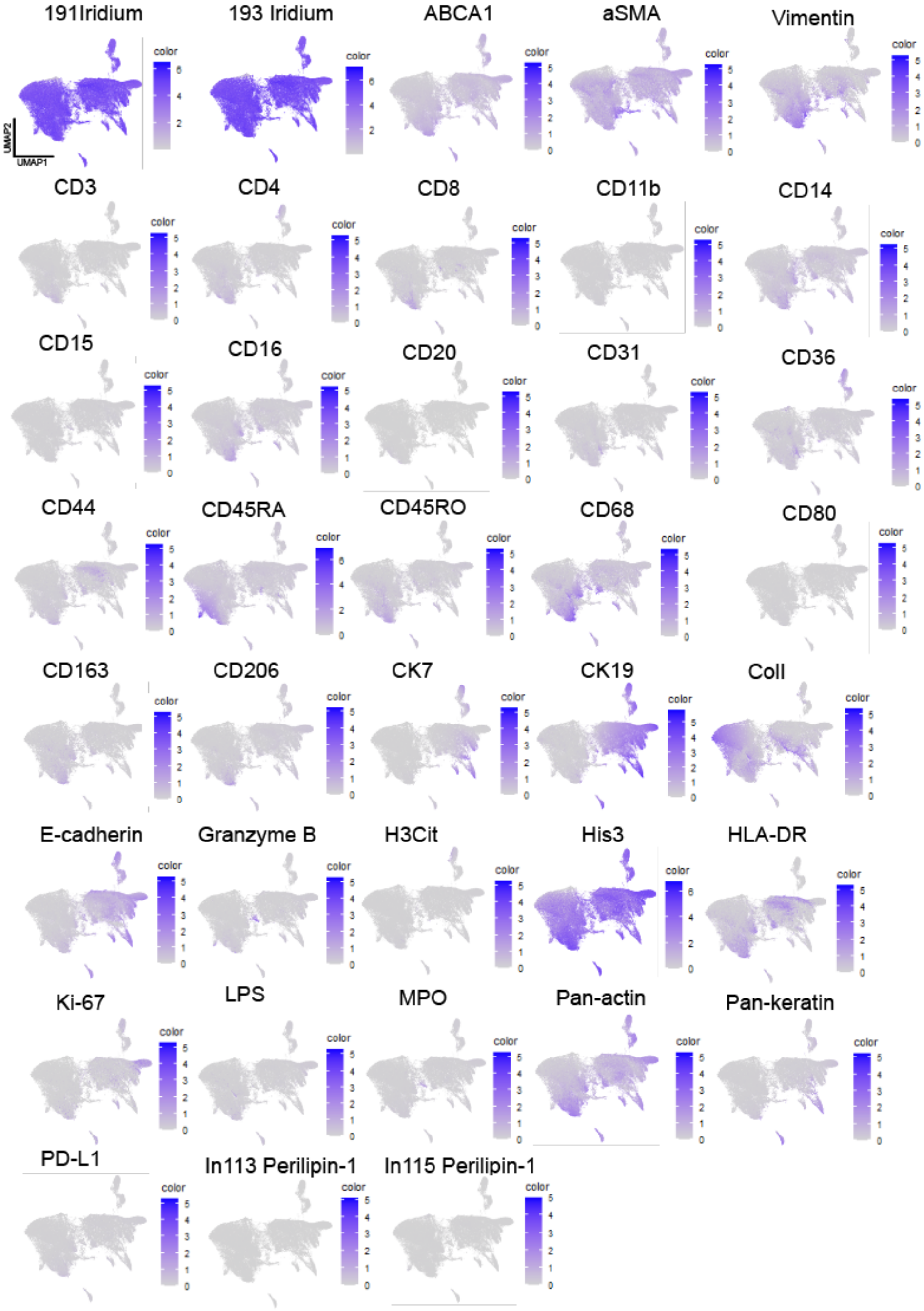
191Iridium
193 Iridium
ABCA1
aSMA
Vimentin
CD3
CD4
CD8
CD11b
CD14
CD15
CD16
CD20
CD31
CD36
CD44
CD45RA
CD45RO
CD68
CD80
CD163
CD206
CK7
CK19
Coll
E-cadherin
Granzyme B
H3Cit
His3
HLA-DR
Ki-67
LPS
MPO
Pan-actin
Pan-keratin
PD-L1
In113 Perilipin-1
In115 Perilipin-1

C

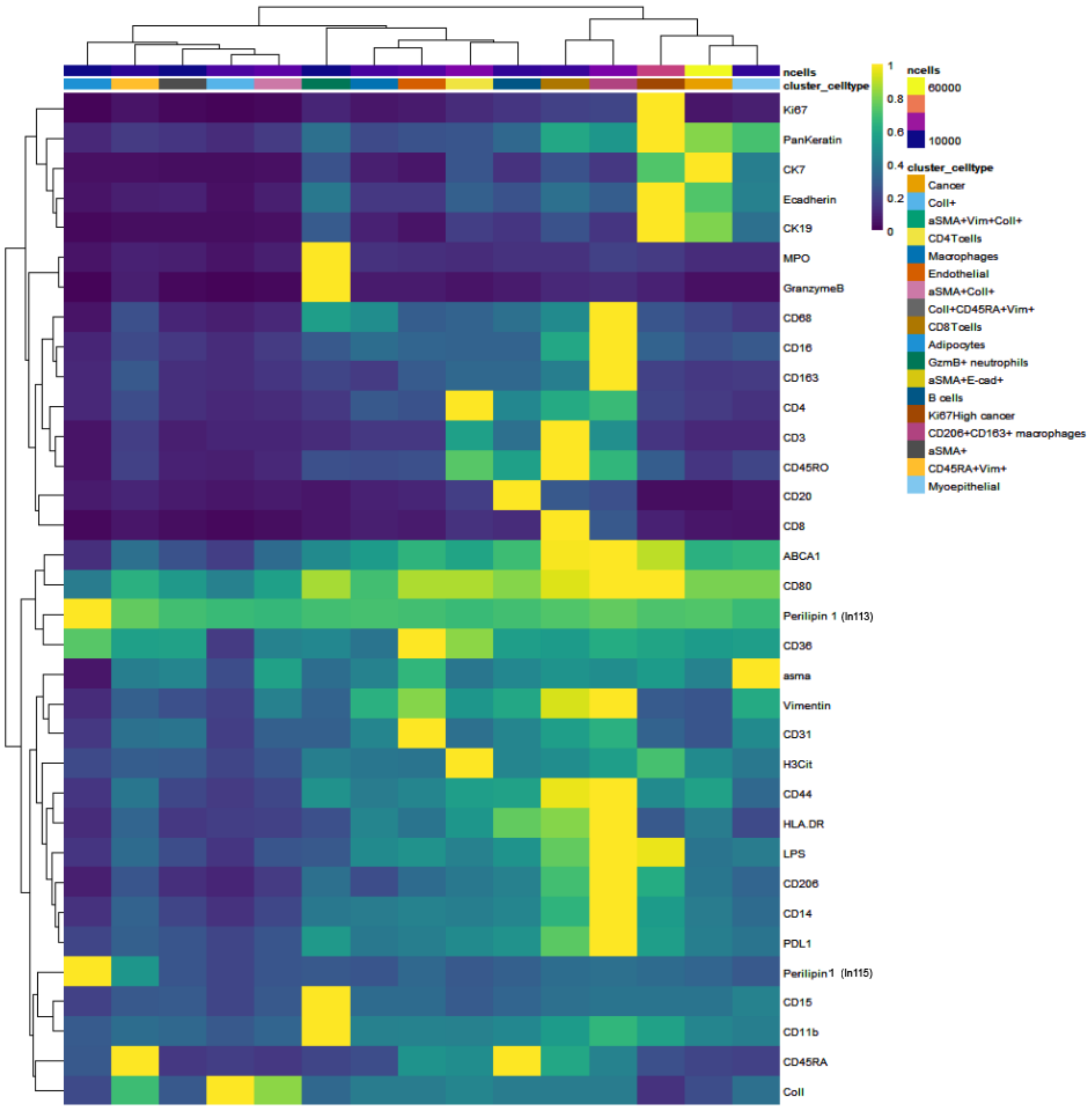

ncells
cluster_celltype
Ki67
PanKeratin
CK7
Ecadherin
CK19
MPO
GranzymeB
CD68
CD16
CD163
CD4
CD3
CD45RO
CD20
CD8
ABCA1
CD80
Perilipin 1 (In113)
CD36
asma
Vimentin
CD31
H3Cit
CD44
HLA.DR
LPS
CD206
CD14
PDL1
Perilipin1 (In115)
CD15
CD11b
CD45RA
ColI
1
0.8
0.6
0.4
0.2
0
ncells
60000
10000
cluster_celltype
Cancer
ColI+
aSMA+Vim+ColI+
CD4Tcells
Macrophages
Endothelial
aSMA+ColI+
ColI+CD45RA+Vim+
CD8Tcells
Adipocytes
GzmB+ neutrophils
aSMA+E-cad+
B cells
Ki67High cancer
CD206+CD163+ macrophages
aSMA+
CD45RA+Vim+
Myoepithelial


D

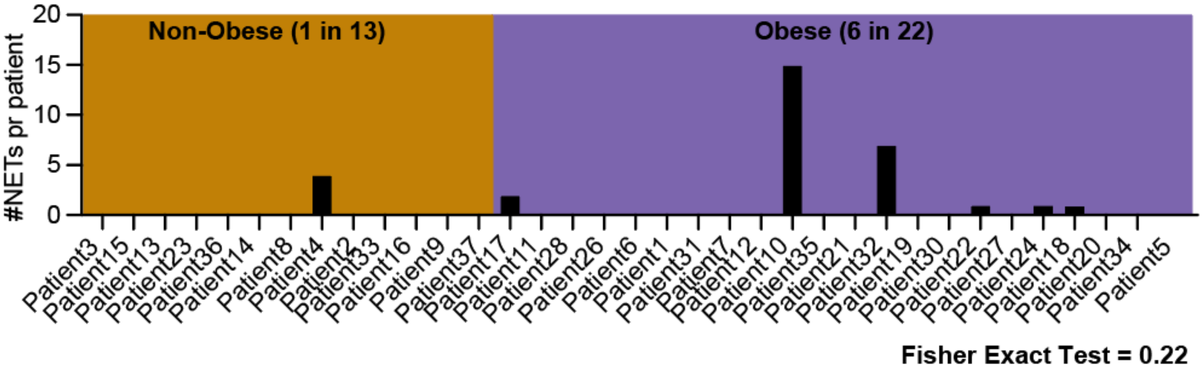

Non-Obese (1 in 13)
Obese (6 in 22)
#NETs pr patient
20
15
10
5
0
Patient3
Patient15
Patient13
Patient23
Patient36
Patient14
Patient8
Patient4
Patient2
Patient33
Patient16
Patient9
Patient37
Patient17
Patient11
Patient28
Patient26
Patient6
Patient1
Patient31
Patient7
Patient12
Patient10
Patient35
Patient21
Patient32
Patient19
Patient30
Patient22
Patient27
Patient24
Patient18
Patient20
Patient34
Patient5
Fisher Exact Test = 0.22

**Fig S3.**

(**A**) Representative image of immunofluorescent staining of $CD68^+$ Cleaved Gasdermin E $(cGSDME)^+$ Cleaved Caspase-3 $(CC3)^+$ cell within the breast TME. CD68 = red, cGSDME = yellow, CC3 = green, and DAPI = blue. Bar plot of the percentage of $CD68^+CC3^+cGSDME^+$ and $CD68^+CC3^+cGSDME^-$ cells within the breast TME among 100 counted $CC3^+CD68^+$ macrophages. (**B**) 2-D UMAP embedding of all cells (both obese ($n$ = 22 patients) and non-obese ($n$ = 13 patients)) in IMC panel 2, colored by arcsinh transformed (cofactor = 1) marker expression. (**C**) Heatmap of the mean marker expression of the cell clusters in IMC panel 2. Cell clusters were generated using PhenoGraph (k = 30). Some clusters were subsequently manually curated. The mean arcsinh transformed (cofactor = 1) marker expressions for each cell cluster are scaled to have a maximum of 1. (**D**) Quantification of NETs (overlap of H3Cit and MPO) within the TME of $PM/ER^-/PR^-$ obese (purple, $n$ = 22) and non-obese (orange, $n$ = 13) patients from the IMC panel 2 data set.

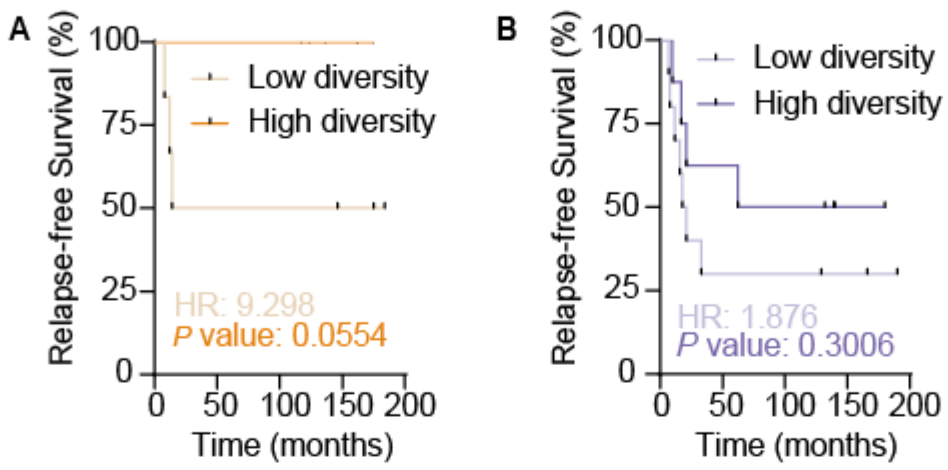


**Figure S4.**

(**A**) Kaplan-Meier plot of Relapse-free Survival of (**A**) non-obese (BMI ≤ 25 kg/m$^2$) and (**B**) obese (BMI > 25 kg/m$^2$) PM/ER$^-$/PR$^-$ breast cancer patients from the in-house data set based on median split by Shannon diversity score of all patients. Patients with high diversity (dark orange, *n* = 6 or dark purple, *n* = 8) had a Shannon diversity score above or equal to median, while patients with low diversity (light orange, *n* = 6, or light purple, *n* = 10) had a Shannon diversity score below median. Log-rank (Mantel-Cox) test *P* value and Hazard ratio (HR) log rank are displayed.

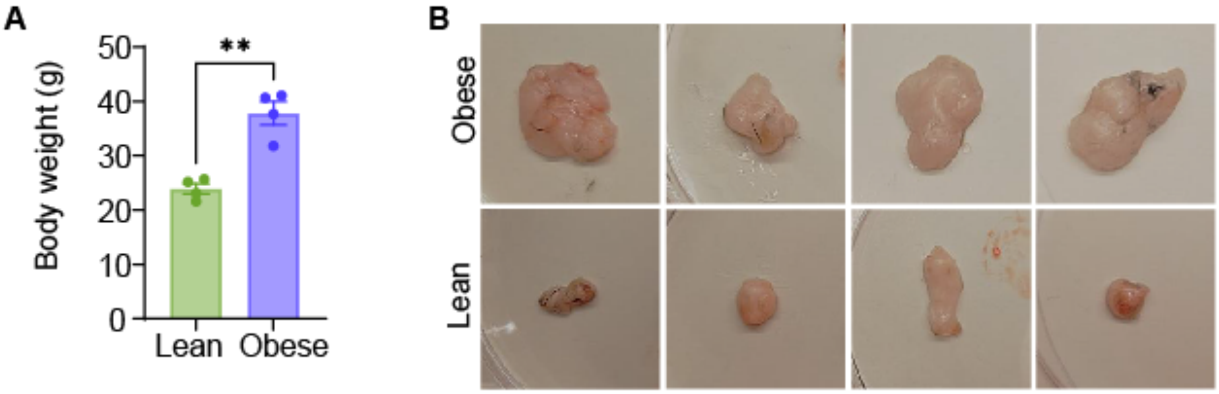


**Figure S5.**

(**A**) Body weight (g) of obese (blue, $n$ = 4) and lean (green, $n$ = 4) mice at day of orthotopic injection of 10 000 EO771 cells. (**B**) Images of the mammary-fat pad (MFP) contralateral to the EO771 tumor from obese ($n$ = 4) and lean ($n$ = 4) mice 22 days after injection.

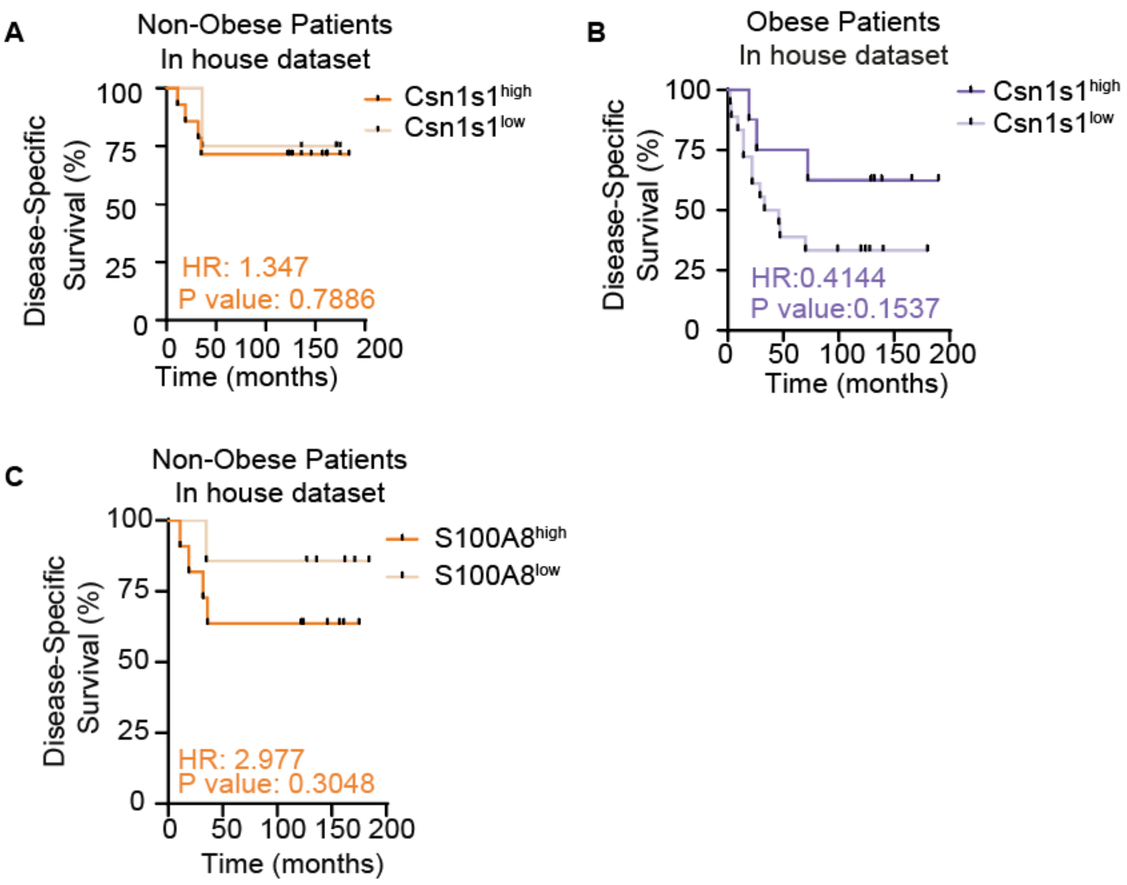


**Figure S6.**

Kaplan-Meier plots of Disease-Specific Survival of (**A**) non-obese (BMI $\leq$ 25) and (**B**) obese (BMI > 25) postmenopausal (> 50 years) $ER^-PR^-$ patients based on tumoral Csn1s1-expression. The median Csn1s1 expression across all included patients was 13.706. Patients were categorized as $Csn1s1^{high}$ (Csn1s1 expression > median) or $Csn1s1^{low}$ (Csn1s1 expression $\leq$ median). Log-rank (Mantel-cox) test *P* value and Hazard ratio (HR) (logrank) are denoted. (**C**) Kaplan-Meier plot of Disease-Specific Survival of non-obese (BMI $\leq$ 25) postmenopausal (> 50 years) $ER^-PR^-$ breast cancer patients from the in-house data set. Patients were separated by median tumoral S100A8 expression (17.572) across all included patients. $S100A8^{high}$ patients had an expression above median, while $S100A8^{low}$ patients had an expression equal to or lower than median. Log-rank (Mantel-cox) test P value and Hazard ratio (HR) (logrank) are displayed.

**Table S1: IMC panel 1**

| Target | Clone | Isotope | Catalog # | Vendor | Concentration/dilutions |
|---|---|---|---|---|---|
| **Perilipin-1** | Polyclonal | 113In/115In | ab3526 | Abcam | 1:200 |
| **Smooth muscle actin** | 1A4 | 141Pr | 3141017D | Fluidigm | 1:200 |
| **H3K4me1** | ERP16597 | 142Nd | ab239402 | Abcam | 1:500 |
| **Vimentin** | D21H3 | 143Nd | 3143027D | Fluidigm | 1:100 |
| **CD14** | EPR3653 | 144Nd | 3144025D | Fluidigm | 1:200 |
| **CD33** | EPR23051-101 | 145Nd | ab269461 | Abcam | 1:100 |
| **CD16** | EPR16784 | 146Nd | 3146020D | Fluidigm | 1:100 |
| **SOX9** | EPR14335 | 147Sm | 3147022D | Fluidigm | 1:100 |
| **Pan-Keratin** | C11 | 148Nd | 3148020D | Fluidigm | 1:100 |
| **CD11b** | EPR1344 | 149Sm | 3149028D | Fluidigm | 1:400 |
| **PD-L1** | E1L3N | 150Nd | 3150031D | Fluidigm | 1:100 |
| **CD31** | EPR3094 | 151Eu | 3151025D | Fluidigm | 1:100 |
| **H3K27me3** | EPR18607 | 152Sm | ab222481 | Abcam | 1:100 |
| **CD44** | IM7 | 153Eu | 3153029D | Fluidigm | 1:200 |
| **CD11c** | Polyclonal | 154Sm | 3154025D | Fluidigm | 1:200 |
| **FoxP3** | 236A/E7 | 155Gd | 3155016D | Fluidigm | 1:100 |
| **CD4** | EPR6855 | 156Gd | 3156033D | Fluidigm | 1:400 |
| **E-Cadherin** | 24E10 | 158Gd | 3158029D | Fluidigm | 1:100 |
| **CD68** | KP1 | 159Tb | 3159035D | Fluidigm | 1:200 |
| **Cytokeratin 19** | EP1580Y | 160Gd | ab195872 | Abcam | 1:100 |
| **CD20** | H1 | 161Dy | 3161029D | Fluidigm | 1:200 |

| | | | | | |
|---|---|---|---|---|---|
| **CD8a** | C8/144B | 162Dy | 3162034D | Fluidigm | 1:200 |
| **CD133** | AC133 | 163Dy | 130-108-062 | Miltenyi Biotec | 1:100 |
| **Cytokeratin 7** | RCK105 | 164Dy | 3164028D | Fluidigm | 1:100 |
| **p-CREB** | 87G3 | 165Ho | 3165034D | Fluidigm | 1:100 |
| **CD45RA** | HI100 | 166Er | 3166028D | Fluidigm | 1:400 |
| **Granzyme B** | EPR20129-217 | 167Er | 3167021D | Fluidigm | 1:200 |
| **Ki-67** | B56 | 168Er | 3168022D | Fluidigm | 1:100 |
| **Collagen type I** | Polyclonal | 169Tm | 3169023D | Fluidigm | 1:400 |
| **CD3** | Polyclonal, C-Terminal | 170Er | 3170019D | Fluidigm | 1:200 |
| **p-ERK1/2** | D13.14.4E | 171Yb | 3171021D | Fluidigm | 1:100 |
| **Caspase-3 cleaved** | 5A1E | 172Yb | 3172027D | Fluidigm | 1:200 |
| **CD45RO** | UCHL1 | 173Yb | 3173016D | Fluidigm | 1:500 |
| **HLA-DR** | YE2/36 HLK | 174Yb | 3174023D | Fluidigm | 1:100 |
| **Pan-Actin** | D18C11 | 175Lu | 3175032D | Fluidigm | 1:100 |
| **Histone H3** | D1H2 | 176Yb | 3176023D | Fluidigm | 1:500 |
| **DNA** | - | 191Ir | 201192B | Fluidigm | 0.3125 µM |
| **DNA** | - | 193Ir | 201192B | Fluidigm | 0.3125 µM |

**Table S2: IMC Panel 2**

| Target | Clone | Isotope | Catalog # | Vendor | Concentration/dilutions |
|---|---|---|---|---|---|
| **Perilipin-1** | Polyclonal | 113In/115In | ab3526 | Abcam | 1:200 |
| **Smooth muscle actin** | 1A4 | 141Pr | 3141017D | Fluidigm | 1:200 |
| **CD15** | W6D3 | 142Nd | 323002 | BioLegend | 1:150 |
| **Vimentin** | D21H3 | 143Nd | 3143027D | Fluidigm | 1:100 |
| **CD14** | EPR3653 | 144Nd | 3144025D | Fluidigm | 1:200 |
| **MPO** | Polyclonal Goat IgG | 145Nd | AF3667 | Biotechne | 1:500 |
| **CD16** | EPR16784 | 146Nd | 3146020D | Fluidigm | 1:100 |
| **CD163** | EDHu-1 | 147Sm | NB110-40686 | Biotechne R&D systems | 1:100 |
| **Pan-Keratin** | C11 | 148Nd | 3148020D | Fluidigm | 1:100 |
| **CD11b** | EPR1344 | 149Sm | 3149028D | Fluidigm | 1:400 |
| **PD-L1** | 73-10 | 150Nd | ab226766 | Abcam | 1:200 |
| **CD31** | EPR3094 | 151Eu | 3151025D | Fluidigm | 1:100 |
| **ABCA1** | Polyclonal | 152Sm | NB400-105 | Novus Biologicals | 1:100 |
| **CD44** | IM7 | 153Eu | 3153029D | Fluidigm | 1:200 |
| **CD80** | Monoclonal Mouse IgG1 Clone # 37711 | 154Sm | MAB140-100 | Biotechne R&D systems | 1:100 |
| **H3Cit** | EPR20358-120 | 155Gd | ab232938 | Abcam | 1:200 |

| | | | | | |
|---|---|---|---|---|---|
| **CD4** | EPR6855 | 156Gd | 3156033D | Fluidigm | 1:400 |
| **CD36** | D8L9T | 157Gd | 39914SF | Cell Signaling Technology | 1:500 |
| **E-Cadherin** | 24E10 | 158Gd | 3158029D | Fluidigm | 1:100 |
| **CD68** | KP1 | 159Tb | 3159035D | Fluidigm | 1:200 |
| **Cytokeratin 19** | EP1580Y | 160Gd | ab195872 | Abcam | 1:100 |
| **CD20** | H1 | 161Dy | 3161029D | Fluidigm | 1:200 |
| **CD8a** | C8/144B | 162Dy | 3162034D | Fluidigm | 1:200 |
| **CD206** | Polyclonal | 163Dy | ab64693 | Abcam | 1:500 |
| **Cytokeratin 7** | RCK105 | 164Dy | 3164028D | Fluidigm | 1:100 |
| **LPS** | WN1 222-5 | 165Ho | HM6011-500UG | Hycultbiotech | 1:500 |
| **CD45RA** | HI100 | 166Er | 3166028D | Fluidigm | 1:400 |
| **Granzyme B** | EPR20129-217 | 167Er | 3167021D | Fluidigm | 1:200 |
| **Ki-67** | B56 | 168Er | 3168022D | Fluidigm | 1:100 |
| **Collagen type I** | Polyclonal | 169Tm | 3169023D | Fluidigm | 1:400 |
| **CD3** | Polyclonal, C-Terminal | 170Er | 3170019D | Fluidigm | 1:200 |
| **Caspase-3 cleaved** | 5A1E | 172Yb | 3172027D | Fluidigm | 1:200 |
| **CD45RO** | UCHL1 | 173Yb | 3173016D | Fluidigm | 1:500 |
| **HLA-DR** | LN3 | 174Yb | 3174025D | Fluidigm | 1:200 |
| **Pan-Actin** | D18C11 | 175Lu | 3175032D | Fluidigm | 1:100 |

| | | | | | |
|---|---|---|---|---|---|
| **Histone H3** | D1H2 | 176Yb | 3176023D | Fluidigm | 1:500 |
| **DNA** | - | 191Ir | 201192B | Fluidigm | 0.3125 μM |
| **DNA** | - | 193Ir | 201192B | Fluidigm | 0.3125 μM |

**Table S3: RT-qPCR Primers**

| Gene | Sequence |
| --- | --- |
| mS100A8_Forward | CAAGGAAATCACCATGCCCTCTA |
| mS100A8_Reverse | ACCATCGCAAGGAACTCCTCGA |
| mJunb_forward | GACCTGCACAAGATGAACCACG |
| mJunb_reverse | ACTGCTGAGGTTGGTGTAGACG |
| mBactin_forward | CATTGCTGACAGGATGCAGAAGG |
| mBactin_reverse | TGCTGGAAGGTGGACAGTGAGG |
